# Molecular mechanisms of long-term light adaptation of an extremophilic alga *Cyanidioschyzon merolae*

**DOI:** 10.1101/2022.03.02.482653

**Authors:** Mateusz Abram, Radek Kaňa, Didrik Olofsson, Filip Pniewski, Barbora Šedivá, Martha Stark, Dylan Fossl, Viktor Slat, Alexander Neumann, Stephen Rader, Joanna Kargul

## Abstract

Oxygenic phototrophs have evolved a remarkable plethora of strategies to react to changes in light intensity and spectral range, which allows them to thrive in a wide range of environmental conditions. Varying light quality and quantity influences the balance between solar energy capture and utilisation in photosynthesis, affecting concomitantly the downstream processes of central carbon and nitrogen metabolism as well as cellular growth and division. Here, we performed a comprehensive analysis of the mechanisms of long-term photoacclimation of an extremophilic red alga *Cyanidioschyzon merolae* that grows in sulphuric hot springs at high temperatures and low pH. By using spectroscopic, confocal fluorescence microscopy, photosynthetic performance measurements and global transcriptome analyses, we identified several molecular mechanisms underlying the long-term adaptation of this acido-thermophilic red alga to varying light intensity and spectral quality. These include: (1) remodelling of the functional antenna size of both photosystems; (2) rearrangement of the PSB/PSII/PSI microdomains within thylakoids; (3) modulation of the photosynthetic performance parameters, especially at the level of non-photochemical quenching, and (4) transcriptional regulation of photosynthesis and its regulatory components as well as downstream metabolic pathways related to ROS detoxification, cell/organelle division, and central carbon and nitrogen metabolism. Such an intricate network of interplay between light-driven reactions and downstream metabolic pathways provides the necessary basis for maintaining the highest photosynthetic performance under light-limiting conditions.

## Introduction

Aquatic eukaryotic algae have evolved for over 1 billion years and are now present in almost every ecological niche on earth. These organisms have adopted strategies to adapt to everchanging light conditions: from fast changes in the irradiation related to weather, through day/night cycles, to seasonal changes in the day length, average irradiation levels, and light penetration of aqueous niches. Eukaryotic algae have subsequently radiated throughout oceanic/fresh water/terrestrial environments, adopting distinctive morphological and developmental strategies for adaptation to diverse light environments. During evolution, algae have adapted to withstand a typical diurnally fluctuating light supply ranging from ~0 to ~2000 μE m^−2^ s^−1^. From the long-term perspective, such extreme changes in light intensity, and variation in light quality, have driven evolution of novel photoreceptors, light harvesting complexes, and photoprotective mechanisms in phototrophic eukaryotes (Archibald and Keeling, 2002; Nowack et al., 2008; Burki et al., 2016; Duanmu et al., 2017).

The photoexcitation of the two main molecular machines of light-dependent reactions of photosynthesis, photosystems II and I (PSII and PSI), must be balanced to ensure the highest efficiency of photochemistry and thus, to optimise the quantum yield of photosynthesis. In low light (LL), the imbalance in the excitation of both photosystems is counteracted by the rapid process of state transitions (ST). The molecular basis of ST differs between prokaryotic cyanobacteria, red algae, and the green lineage (including green algae and higher plants). In the latter group of phototrophs, the ST process involves the reversible phosphorylation/dephosphorylation of the light harvesting components of PSII and the distinct LHCII subunits. The corresponding kinases and phosphatases have been identified through mutagenesis studies (Kargul and Barber, 2008; Wei et al., 2015; Bolychevtseva et al., 2021). The phosphorylated LHCII complexes undergo a conformational change followed by their subsequent detachment from the PSII complex and attachment to the specific domain of the PSI core by molecular recognition (Bos et al., 2017; Kosuge et al., 2018). The signalling components include the LHCII kinase STN7 (Depège et al., 2003; Bellafiore et al., 2005; Pesaresi et al., 2009) and the PPH1/TAP38 phosphatase (Pribil et al., 2010; Shapiguzov et al., 2010).

In the case of cyanobacteria and red algae, LHCII antenna complex is absent and substituted with a huge (over 6 MDa in size) phycobilisome complex (PBS) that serves as the peripheral antenna to photosystems. The PBS complex efficiently harvests blue/green light due to its phycobilin pigment composition and its unique structural organisation of rod, core, and linker components (Yamanaka et al., 1980; MacColl, 1998; Kerfeld et al., 2003; Zhang et al., 2017; Sauer et al., 2021). In PBS-containing phototrophs, four non-exclusive models of ST have been proposed: (1) model with the mobile PBS antenna; (2) excitation energy spill-over model; (3) PBS detachment model; and (4) model of quenching in the PSII reaction centre (Fujita et al., 1994; Yokono et al., 2011; Calzadilla and Kirilovsky, 2020). The *mobile antenna* model explains the regulation process as analogous to the LHCII movement between both photosystems in the green lineage. However, it needs to be noted that the actual importance of this model as an *in vivo* mechanism in cyanobacteria and red algae is still rather questionable especially in light of photophysics effects during FRAP measurements used for observation of the putative PBS movement (see discussion in Calzadilla and Kirilovsky, 2020). The second *spill-over* mechanism requires the specific arrangement of the photosynthetic complexes in the thylakoid membrane, which allows for the energy transfer between both photosystems via PBS (McConnell et al., 2002). In cyanobacteria and red algae, PSI and PSII complexes are positioned in very close proximity, in accordance with the concept of photosynthetic supermegacomplex formation, and bearing in mind that the thylakoid membrane is not structured into the stacked and lamellar regions present in the green lineage (Cunningham et al., 1989; Rakhimberdieva et al., 2001; Kaňa et al., 2014). The last two mechanisms (PBS decoupling model and RC quenching model) were proposed more recently; they suggest that the fluorescence changes visible during ST could be caused by a transient PBS de-coupling from photosystems or by the PSII RC-based quenching mechanisms. Both these mechanisms then result in fine-tuning of photosynthetic electron flow with PBS complex and PSII core complex acting as the central regulatory unit during ST (Delphin et al., 1996; Krupnik et al, 2013; Ueno et al., 2017; Zlenko et al., 2017).

The short-term acclimation process triggered in HL in the “green” model organisms (higher plants and green algae) corresponds to a rapidly reversed non-photochemical quenching (NPQ) process (qE component) which involves excess energy dissipation as heat through pH controlled process localised in the peripheral or inner antennae of PSII (Baker, 2008; Goss and Lepetit, 2015). However, in PBS-containing organisms, i.e., cyanobacteria and even more in red algae, the pH control of NPQ is not so clear (compare Delphin et al., 1995; Delphin et al., 1998 and Krupnik et al., 2013). Additionally, the NPQ mechanisms in cyanobacterial PBS antenna, clearly occurs by a pH-independent mechanism through conformational change in the PBS complexes triggered by the blue light sensor, the Orange Carotenoid-binding Protein (OCP) (Campbell et al., 1998; Kirilovsky, 2007; Sonani et al., 2018; Dominguez-Martin et al., 2021). In higher plants and green algae, the NPQ mechanism includes also the pH-dependent xanthophyll cycle activity (Yamamoto et al., 1962; Hager, 1967), and an involvement of a pH-sensitive protein known as PsbS (Li et al., 2002). In cyanobacteria and in red algae, both these pH-dependent mechanisms are absent as the PsbS protein and xanthophyll cycle do not exist in these organisms. Additionally, the OCP protein important in cyanobacteria is absent in the red algae genome, therefore the precise mechanism of NPQ is based on another mechanism represented probably by reversible NPQ photoprotection mechanism directly in the RC of PSII (Krupnik et al., 2013).

The long-term adaptation to fluctuating light quantity and spectral quality has been shown to result in significant alteration of the composition and organisation of the thylakoid membrane. In LL-grown higher plants, chloroplasts are characterised by increased thylakoid membrane stacking and higher ratios of LHCII/PSII and PSI/PSII proteins (Dietzel et al., 2008). In contrast, HL stressed plants showed the opposite effects together with elevated abundance of ATP synthase and cytochrome *b*6*f* (cyt *b*6*f*) complexes relative to total the chlorophyll (Chl) content (Walters and Horton, 1995; Bailey et al., 2001; Tikkanen et al., 2006; Ballottari et al., 2007; Wientjes et al., 2013a; Wientjes et al., 2013b; Ware et al., 2015; Schumann et al., 2017; Vialet-Chabrand et al., 2017; Flannery et al., 2021). Consequently, HL-grown plants have a higher capacity for linear and cyclic electron transfer and CO2 assimilation, coupled with an increased resistance to photoinhibition. On the contrary, LL-acclimated plants utilise low irradiance more efficiently (Boardman, 1977; Anderson et al., 1988; Gray et al., 1996).

In contrast to higher plants, there have been only few studies on photoadaptation mechanisms in PBS-containing red algae, and even less in extremophilic microalgae. However, the remarkable robustness of photosynthesis in these phototrophs is clear as they can cope with conditions that are hostile to most other organisms. The extremophilic red microalgae, Cyanidiales, are in fact the only eukaryotic photosynthetic organisms thriving in hot, volcanic, sulphur-rich springs, which are not only highly acidic (pH 0.05-4), but are also rich in heavy metals such as cadmium, nickel, iron, and arsenic (Seckbach, 1994; Ciniglia et al., 2004). Recent bioinformatic analysis addresses the origin and position of the red algae in the phylogenetic tree (Miyagishima and Tanaka, 2021; Sato, 2021). Due to the extensive analysis of the increasing number of the fully annotated genomes and the recent discovery of the heterotrophic eukaryotic phylum Rhodelphidia, recently a new proposal has been put forward regarding the evolutionary origin of red algae (Burki et al., 2016; Lee et al., 2017; Gawryluk et al., 2019) questioning the previous hypotheses (Seckbach, 1994; Seckbach and Chapman, 2010). It is the so-called the rhodoplex hypothesis proposing that the red algae plastid emerged during evolution through a series of independent endosymbiosis events (Miyagishima and Tanaka, 2021; Strassert et al., 2021). The latter hypothesis could explain the chimeric nature of the photosynthetic apparatus in Cyanidiales, as well as the simplicity of the genomic organisation in these algae (Kuroiwa, 1998; Miyagishima and Tanaka, 2021).

One of the six species encountered among the Cyanidiales is *Cyanidioschyzon merolae*, a unicellular red alga discovered in the biomats present in the volcanic area *Campi Flegrei*, Italy. The natural environment of this alga is characterised by moderately high temperatures (40-57 °C) and very low pH (pH 0.2-4) due to the high concentration of the sulphuric acid (Luca et al., 1978). *C. merolae* is an obligate photoautotroph with a small genome and simple subcellular organisation, as it contains a single nucleus, a single mitochondrion, and a single chloroplast (Luca et al., 1978; Kuroiwa, 1998). Contrary to other Cyanidiales species, and in contrast to the green lineage, *C. merolae* cells do not possess a cell wall or vacuoles. Instead, a set of small polyphosphate vacuoles was discovered together with starch inclusions (Yagisawa et al., 2007; Yagisawa et al., 2009). Contrary to higher plant chloroplasts, the *C. merolae* plastid structure is based on simple multiple layers of thylakoid membranes without specialised structures, such as the grana that are present in green algal and higher plant chloroplasts (Ohashi et al., 2010; Allen et al., 2011).

Due to its simple cellular ultrastructure and genome organisation, *C. merolae* has been used as a model organism to study the evolution and molecular mechanisms of the cell cycle, cell signalling, and circadian rhythm processes (Miyagishima and Tanaka, 2021). It is also becoming an important model organism for the study of the evolution and mechanism of pre-mRNA splicing (Stark et al., 2015; Black et al., 2016; Reimer et al., 2017; Garside et al., 2019). More recently, much attention has been devoted to deciphering the molecular mechanisms of photoadaptation in this extremophile, specifically by dissecting the structure and function of the photosynthetic molecular machinery that is closely related to that of other eukaryotic phototrophs, albeit with significant exceptions. The photosystems with associated antenna proteins of *C. merolae* are chimeric in nature having both cyanobacterial and green lineage structural traits, in line with the above-mentioned rhodoplex hypothesis (Miyagishima and Tanaka, 2021). While the PSI complex bears a strong resemblance to plant counterparts, although it contains exclusively chlorophyll *a* (Chl*a*) molecules (Busch et al., 2010; Haniewicz et al., 2018; Pi et al., 2018; Antoshvili et al., 2019), PSII is similar to the cyanobacterial counterpart (Krupnik et al., 2013; Ago et al., 2016), specifically in having the cyanobacterial-like PBS complex functioning as a peripheral antenna. In addition, its pigment composition is similar to its cyanobacterial counterpart in its lack of phycoerythrin and presence of phycocyanin (PC) and allophycocyanin phycobilins (Cunningham et al., 2007; Zlenko et al., 2017). The LHCII complex is absent in this extremophilic alga.

Other cyanobacterial traits present in *C. merolae* are the presence of the PsbV and PsbU subunits of the OEC and the unique fourth subunits of the OEC, the 20-kDa PsbQ’ protein that is distantly related to the green algal and higher plant counterparts (Enami et al., 2008; Ifuku, 2015). The latter protein has been identified through structural studies of a red algal PSII complex in the vicinity of the inner antenna CP43 protein, close to the membrane plane (Krupnik et al., 2013; Ago et al., 2016). The function of PsbQ’ is to facilitate binding of the crucial oxygen evolving complex (OEC) subunit, PsbO (Enami et al., 1998). Interestingly, despite the structural similarity of red algal PSII to its cyanobacterial counterpart (with a notable presence of the eukaryotic PsbW subunit in the red algal PSII), the water exchange kinetics of the *C. merolae* OEC have been shown to be significantly different from those of other studied organisms and point to minor differences in the mechanism of water oxidation (Nilsson et al., 2014; Pham et al., 2019).

Recently, we together with other groups have identified several distinct and unique molecular mechanisms underlying the unprecedented robustness of the *C. merolae* photosynthetic apparatus to extreme conditions, especially in variable light. The *first* mechanism is based on the accumulation of a well-known photoprotective carotenoid, zeaxanthin; it has been localised in both the peripheral light harvesting antenna of PSI (LHCI), as well as in the PSII and PSI reaction centres upon exposure to HL (Krupnik et al., 2013; Tian et al., 2017; Haniewicz et al., 2018; Pi et al., 2018; Antoshvili et al., 2019). The specific pigment molecules identified in the recent X-ray and cryo-EM structures of *C. merolae* PSI complex were proposed to form discrete energy transfer pathways in moderate light (ML) (Pi et al., 2018; Antoshvili et al., 2019). The *second* mechanism involves the structural and functional remodelling of the LHCI antenna complex within the PSI-LHCI supercomplex. It is shown by the presence of the two PSI-LHCI isomers whose stoichiometric ratio alters in response to the dynamic changes in illumination (Tian et al., 2017; Haniewicz et al., 2018). The structural remodelling of the Lhcr subunits forming the LHCI antenna is additionally accompanied by the dissociation of the PsaK core subunit of PSI, especially under HL conditions (Haniewicz et al., 2018). The *third* mechanism is the accumulation of low-energy (red) chlorophyll molecules, mainly in the LHCI subunits under light-limiting conditions, that exert a dual function: as the light harvesting molecules in LL or as energy traps under HL conditions (Abram et al., 2020). Moreover, in HL the LHCI antenna appears functionally decoupled from the PSI reaction centre, as shown by the recent time-resolved spectroscopic studies on isolated PSI complexes (Abram et al., 2020; Chang et al., 2020). Finally, the *fourth* mechanism of photoadaptation is based on the discovery of the reversible RC-based NPQ in the *C. merolae* PSII reaction centre that is triggered by acidification of the thylakoid lumen upon exposure to HL saturating pulses (Krupnik et al., 2013). All the above characteristics are likely to provide the basis for the remarkable robustness and sustained high photochemical activities of the *C. merolae* photosynthetic apparatus, as observed across a wide range of light, temperature, and pH conditions (Krupnik et al., 2013; Haniewicz et al., 2018).

Here, we have investigated the mechanisms of long-term photoacclimation of *C. merolae* cells by spectroscopic and confocal fluorescence microscopy analyses in conjunction with monitoring changes of photosynthetic performance and global transcriptome regulation. We show that the photoadaptation mechanisms to varying light quantity and quality involve: (1) alteration of the effective antenna size of both photosystems; (2) induction of two distinct NPQ responses in eHL versus LL and ML cells, (3) formation and dynamic remodelling of the photosynthetic microdomains in the thylakoid membranes, and (4) remodelling of the transcriptional regulation of photosynthesis and downstream metabolic pathways related to ROS detoxification, cell/organelle division, as well as central carbon and nitrogen metabolism.

## Results

### Remodelling of the light harvesting antennae and photosystem stoichiometry at various light regimes

The global changes in the absorption of the intact cells were examined at varying light quantity (light intensity, Figure 1A) and spectral light quality (Figure 1B and 1C). The various intensities of white light regimes (LL, ML and eHL; Figure 1A) during growth resulted in the five characteristic peaks in the absorption spectra representing the three prominent pigments, chlorophyll a (Chl*a*), phycobilins embedded in the phycobilisome complex (PBS) and carotenoids (Car). The Soret band (410–440 nm) and Q*y* band (670–680 nm) correspond to Chl*a* absorption, the PBS band (600–640 nm) represents absorption of phycocyanin (PC); Car are typically detected in a 470-500 nm range by a composite absorption of several bands (Ueno et al., 2015). Upon normalisation of the absorption spectra at the Q*y* band, we observed a slight red shift of the Q*y* peak in LL-grown cells (680.5 nm) and eHL-adapted cells (680 nm) compared to the Q*y* peak of ML-adapted control (679.5 nm) likely due the accumulation of the so-called red Chl pigments. These serve as intermediate energy traps for promoting uphill energy transfer from the peripheral to the bulk and/or inner core Chl molecules and ultimately to the P700 RC of the eukaryotic and cyanobacterial PSI (Croce et al., 2000; Gobets and van Grondelle, 2001; Ihalainen et al., 2002; Jennings et al., 2003; Gibasiewicz et al., 2005; Melkozernov et al., 2005; Engelmann et al., 2006; Snellenburg et al., 2013; Abram et al., 2020). The highest PBS/Chl*a* (680 nm) ratio was observed for the LL cells, whereas eHL cells showed the lowest PBS/Chl*a* (680 nm) ratio. This indicates a relative decrease of the PBS content with the increase of white light illumination during cell cultivation. The opposite effect was observed for the Car peaks which were significantly higher in eHL cells compared to the LL and ML counterparts.

**Figure 1.**
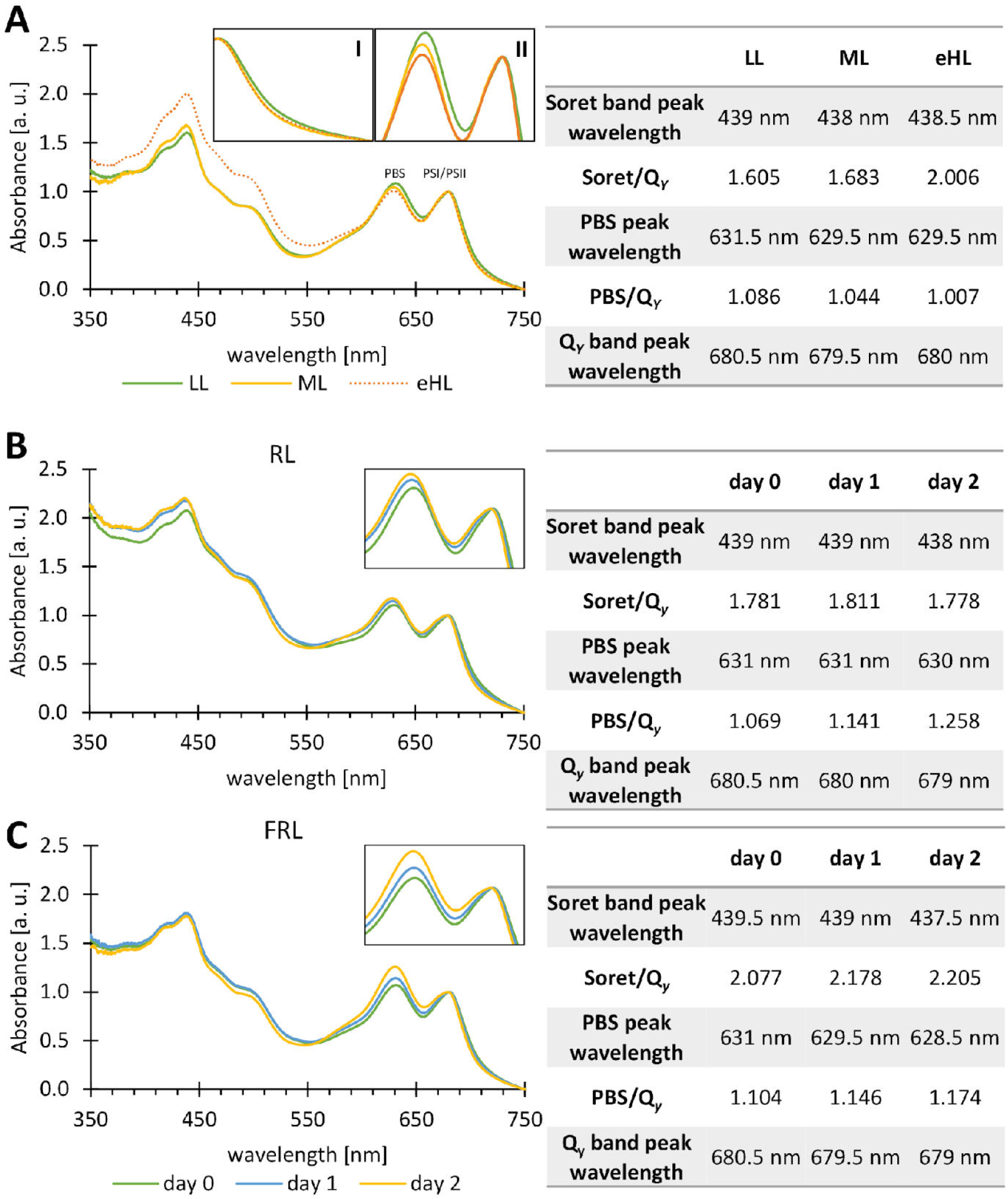
Absorption spectra of the *C. merolae* cultures under various light intensity and spectral quality. Absorption spectra of the cultures grown under different light regimes (LL, ML, eHL, RL and FRL) for up to 48 h. Ratio and peak wavelength are shown in the corresponding tables. Spectra were obtained from two cultures grown at the same time (n=2) and averaged. **(A)** Absorption spectra of the cells grown at different white light intensities: LL (35 μmol photons m^−2^ s^−1^), ML (90 μmol photons m^−2^ s^−1^) and eHL (350 μmol photons m^−2^ s^−1^) PAR, as described in Haniewicz et al. (2019). *Inset I:* close-up of 678-750 nm range; *Inset II:* close-up of 600-700 nm range **(B)** Absorption spectra of the cultures grown under continuous RL illumination. **(C)** Absorption spectra of the cultures grown under continuous FRL irradiance. Spectra were normalised at a Q_*y*_ band.

Prolonged exposure of cells to varying light quality showed different spectral effects. Adaptation of cells to RL or FRL (48 h) induced significant changes, namely 1.5 nm blue shift of the Q*y* Chl*a* band position as well as 6% and 18% increase of the relative absorbance of the PBS normalised to that of Chl*a* (680 nm) (Figure 1B, C). However, no significant changes were detected in the relative Car content under long-term RL and FRL cell adaptation.

The remodelling of the effective PSI and PSII antenna size was analysed by measuring 77K fluorescence of the *C. merolae* cells adapted to growth under various illumination conditions upon Chl*a* excitation at 435 nm (Figure 2). Firstly, we examined changes in the antenna size during long-term adaptation to varying white light intensity (Figure 2). When emission peaks were normalised to the PSII emission peak (685 nm), an over 2-fold increase of the PSI emission peak (728.5 nm) was observed for LL-adapted cells compared to ML control (728 nm), indicating a significant increase in the PSI antenna during long-term LL adaptation. Conversely, the PSI emission in eHL cells decreased to 79% compared to PSI emission in ML cells upon increasing the light intensity (Figure 2A). As the relative content of PBS is slightly lower in eHL conditions and higher during LL adaptation, the changes of the PSI emission peak intensity may reflect changes in the PSII/PSI stoichiometry, namely an increase in the relative content of PSII compared to PSI complexes and/or energy transfer from PBS to PSII upon increasing white light intensity. These changes may also reflect energy spill-over from PSII to PSI via PBS antenna, especially under LL conditions. The latter conclusion is supported by the increase of the PBS emission peak in the excitation spectrum recorded for LL-adapted cells at PSI absorption maximum (728 nm) (Figure 2B). Interestingly, the maximum of PSI emission was red-shifted by 0.5 nm in LL cells (728.5 nm) and blue-shifted by 7 nm in eHL cells (721.5 nm) compared to PSI emission in ML cells peaking at 728 nm (Figure 2A). This observation may correspond to structural changes in LL/eHL PSI complexes with regards to the functional antenna association.

**Figure 2.**
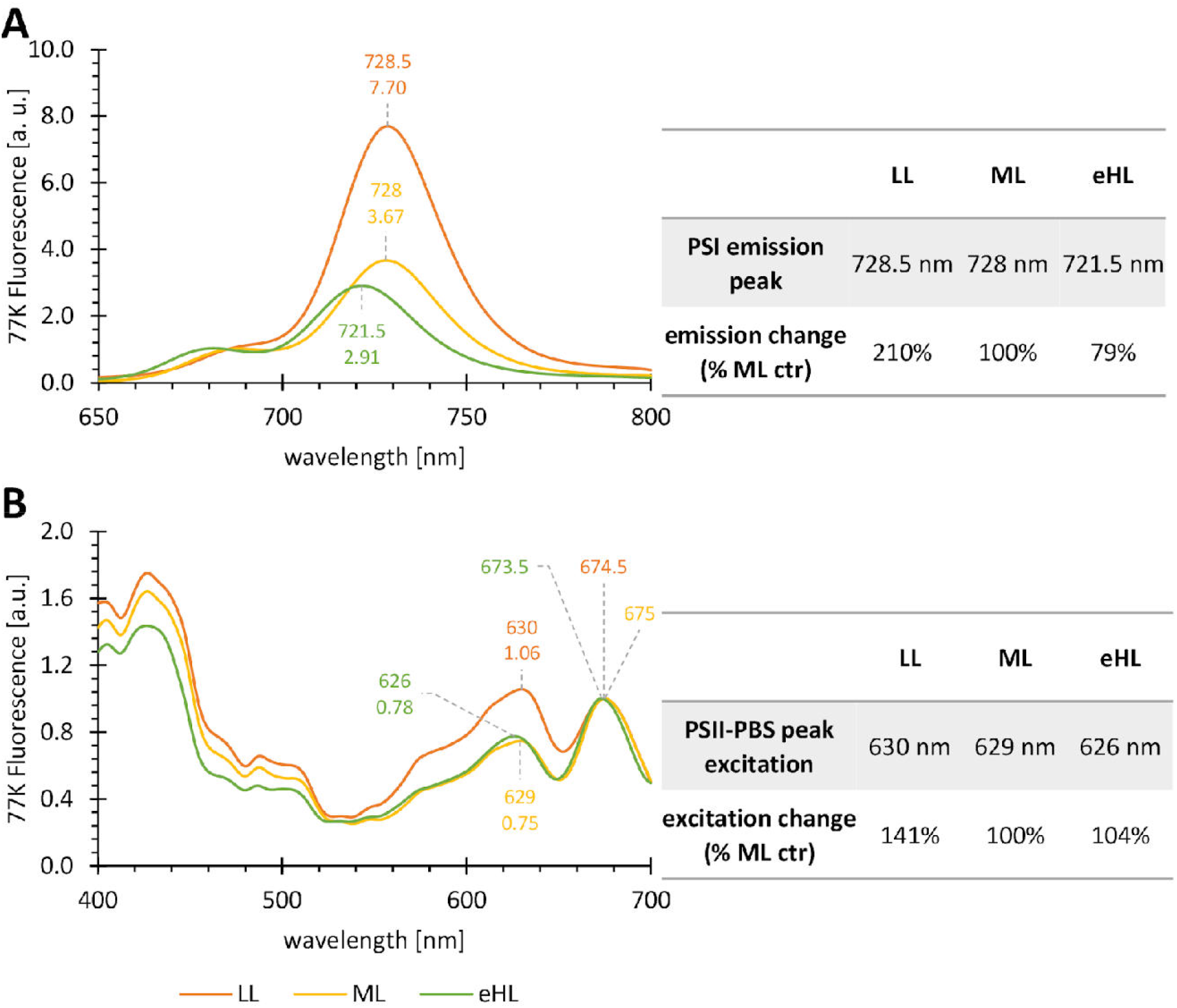
77K fluorescence from cells exposed to various white light regimes. **(A)** Emission spectra of cells obtained by Chl*a* excitation at 435 nm and normalised at 685 nm (PSII emission). PSI emission maxima and PSI/PSII peak ratios are marked above the spectra. Changes in the emission maxima are shown on the right. **(B)** Excitation spectra obtained following excitation of cells at 400 to 700 nm and recording emission at 728 nm (PSI). Spectra were normalised to the PSI excitation maximum (673.5-675 nm). Peak wavelengths and the peak ratio of PSII-PBS/PSI is displayed above the spectra. Changes in the PSII-PBS excitation peak ratio is displayed on the right. Ctr, control.

In the next step, we investigated the putative changes to the effective antenna during adaptation to illumination of the cells with varying spectral quality (Figure 3). In RL- and FRL-adapted cells, 77K PSI emission maxima were blue-shifted by 3 and 5 nm, respectively during the 48 h acclimation (Figure 3A), which may indicate decoupling of the effective PSI antenna subpool, possibly in the process of ST and/or NPQ (see below). In support of the latter hypothesis, upon normalisation of the emission spectra to the PSII peak (685 nm) the effective PSI antenna decreased to 41% of the control (t= 0 h in Fig. 3A) in RL cells. Similarly, FRL treatment decreased the PSI antenna size down to 58% of the control (t=0 in Figure 3B). These changes in the effective antenna size of PSI are may reflect modulation of the PSII/PSI stoichiometry, i.e., an increase in the relative content of PSII complex compared to PSI, during long-term RL and FRL adaptation of the *C. merolae* cells. Both RL and FRL treatments resulted in lower PBS emission in the excitation spectra, which may be due to increased energy transfer from PBS to PSII during RL/FRL long-term adaptation of the *C. merolae* cells.

**Figure 3.**
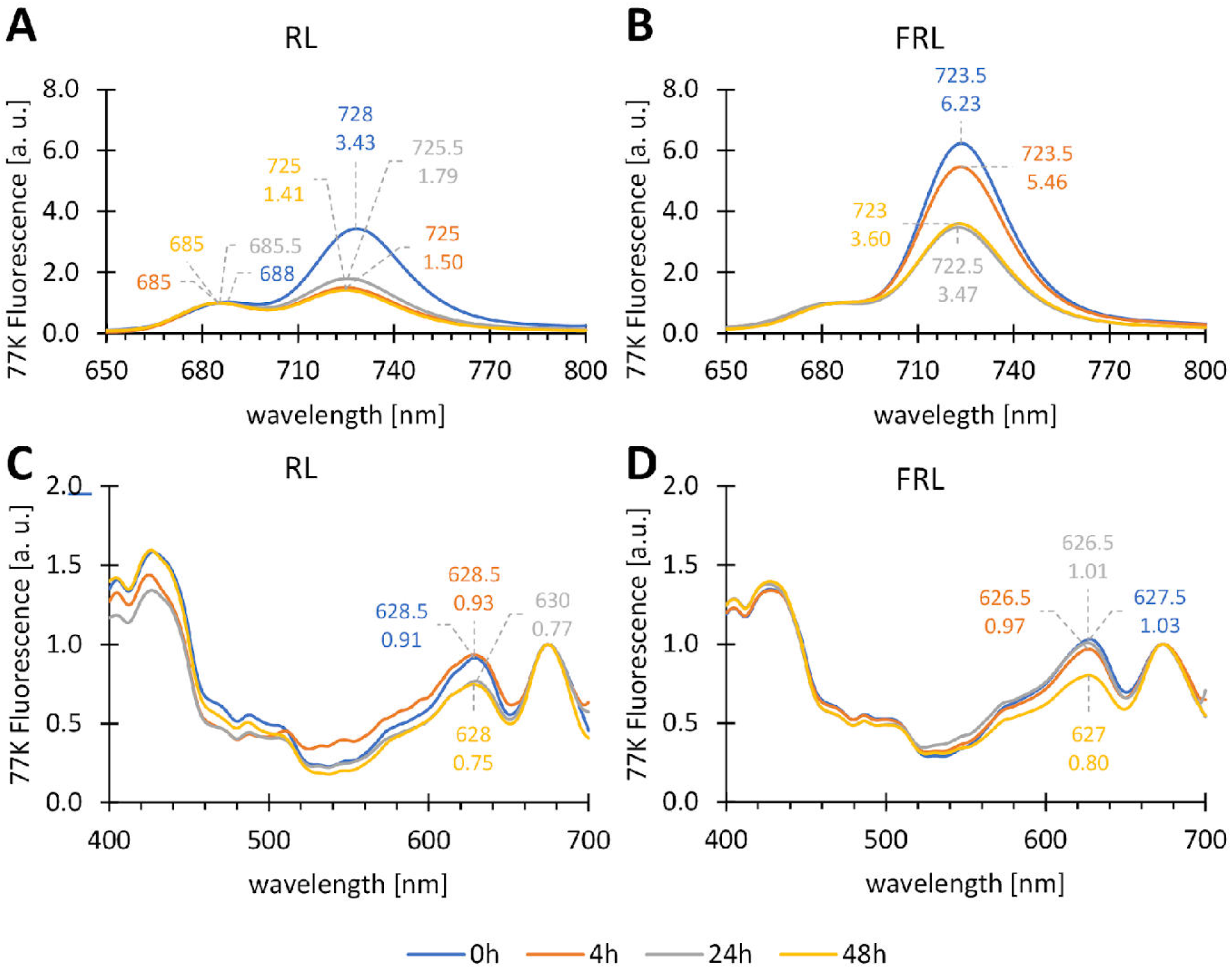
77K fluorescence from cells exposed to illumination of varying spectral quality. Steady state fluorescence spectra obtained from the cells at specific time points: 0h, immediately prior to illumination conditions change and after 4, 24 and 48 h of continuous RL or FRL illumination. **(A, B)** Emission spectra obtained by Chl*a* excitation at 435 nm normalised at 685 nm (PSII emission). PSI emission maxima and PSI/PSII peak ratios are marked above the spectra. **(A)** PSII emission maxima were also displayed, in **(B)** PSI fluorescence overlapped PSII fluorescence. **(C, D)** Excitation spectra obtained following excitation of cells at 400-700 nm and recording emission at 728 nm (PSI). Spectra were normalised to the PSI/PSII peak (673.5-675 nm). Peak wavelengths and the peak ratios of PSII-PBS/PSI are displayed above the spectra. **(A, C)** RL-grown cells. **(B, D)** FRL-grown cells. For peak wavelengths and peak ratios, see **Table 1**.

### Changes of photosynthetic performance at various light intensity and spectral quality

In order to probe potential changes in the photosynthetic performance during long-term adaptation of the *C. merolae* cells to various light conditions, we measured the main photosynthetic parameters of PSII *in vivo* (Baker, 2008). We examined three main parameters related to PSII photochemical performance and non-photochemical protective response. The maximal quantum efficiency of PSII (F_v_/F_m_) reached the highest value for ML-adapted cells and slightly decreased (by 10%) in LL and eHL cells (Figure 4A). The F_v_/F_m_ ratio was not affected by the quality of light and generally remained stable after transition to both RL and FRL conditions (Figure 4C). The process of photoacclimation to light quality/quantity was better visible in the actual PSII efficiency in light (ΦPSII); this parameter was significantly lower in LL cells compared to ML and eHL counterparts (Figure 4B). Simultaneously, the ΦPSII parameter was lower in FRL conditions, and increased to 167% of the control (white light-adapted cells) during RL adaptation of the cells (compare t=0 and t=48 h in Figure 4D).

**Figure 4.**
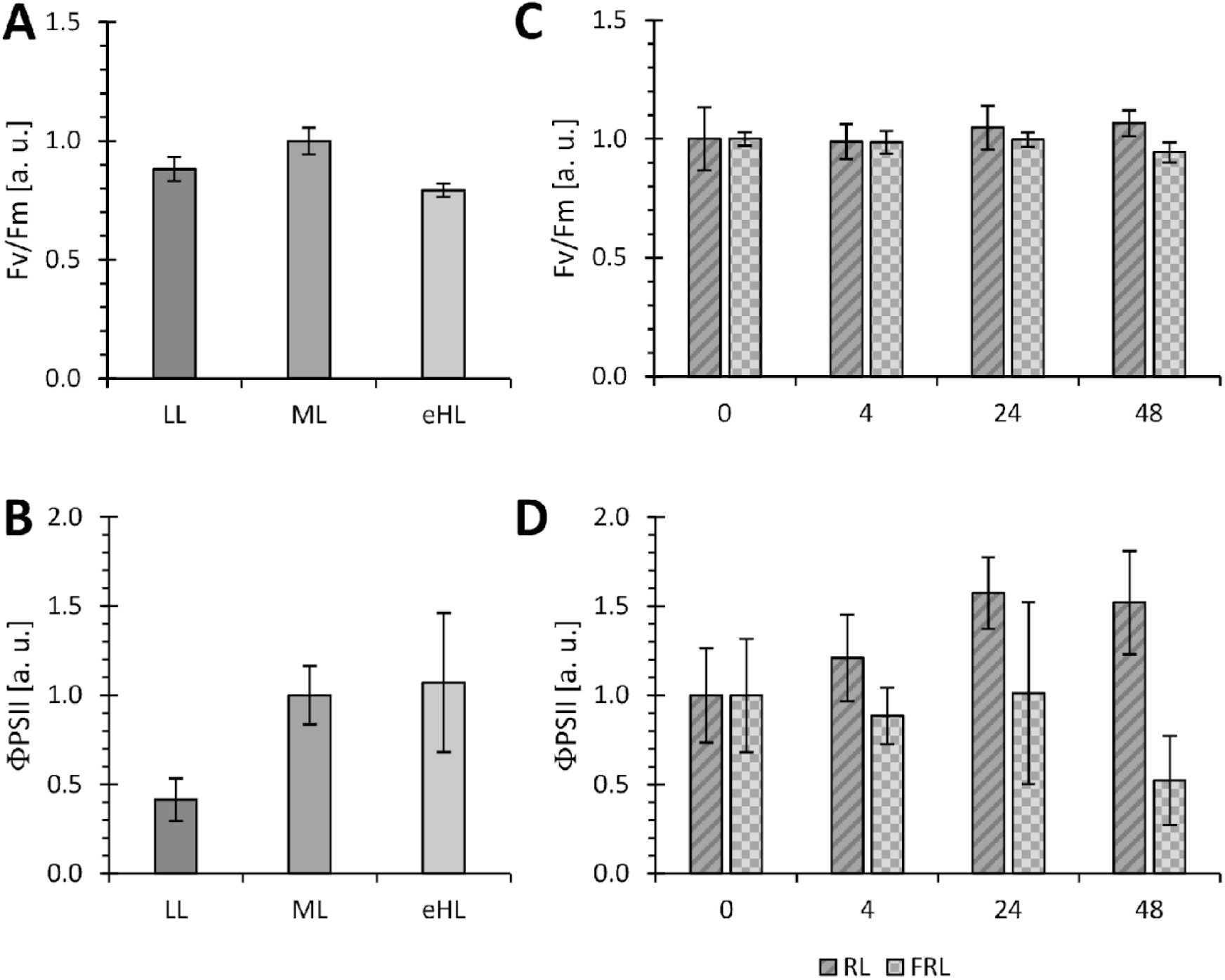
Photosynthetic performance of cells exposed to various light quantity and quality. PAM measurements were carried out for the same time points of cell culture growth curves as for the steady state 77K fluorescence measurements. **(A, B),** photosynthetic parameters obtained from whole cells grown at various white light intensities for up to 48 h. **(C, D)** photosynthetic parameters obtained from the whole cells exposed to RL and FRL illumination for up to 48 h. Each value is the mean (± SD) of 3 technical replicates obtained from 2 independent biological samples (n=2).

The important difference in long-term acclimation of the *C. merolae* cells to various light regimes was strikingly visible through the kinetic analysis of the photoprotective response, non-photochemical quenching, NPQ (Figure 5), by monitoring quenching of the maximal Chl*a* fluorescence of PSII (F_m_). To this end, cells were initially exposed to low blue light (intensity 81 μmol photons m^−2^ s^−1^, λ = 460 nm) and then high-intensity blue light (1374 μmol photons m^−2^ s^−1^, λ = 460 nm), both accompanied by high-intensity red flashes (4000 μmol photons m^−2^ s^−1^, duration 400 ms). The protocol followed the standard procedure used for PBS-containing cyanobacteria (Kirilovsky et al., 2014), whereby the first low-blue light illumination period is applied to induce State 2-to-State 1 transition (Canonico et al., 2021). Surprisingly, we did not observe a typical state transition behaviour (except for eHL-adapted cells) in the ‘low light’ phase of illumination. Instead, we observed two distinct NPQ phenomena, low blue light effect of F_m_ decrease to F_m1_’ (see the first “gray” period in Figure 5) and effect of high-intensity blue light (see the further decrease from F_m1_’ to F_m2_’ in Figure 5). The first multi-turnover flash-induced phenomenon was observed clearly only in LL- and ML-adapted cells, which showed quenching of dark-adapted maximal F_m_ value in low intensity blue light (Figure 5A, B). Additionally, in these cells high-intensity blue light-induced the NPQ response (see decrease from F_m1_’ to F_m2_’ in Figure 5). The data suggest that the drop in F_m1_’ was not caused by low blue light but rather it was the effect of flash-induced NPQ that was previously observed in red algae (Delphin et al., 1998).

**Figure 5.**
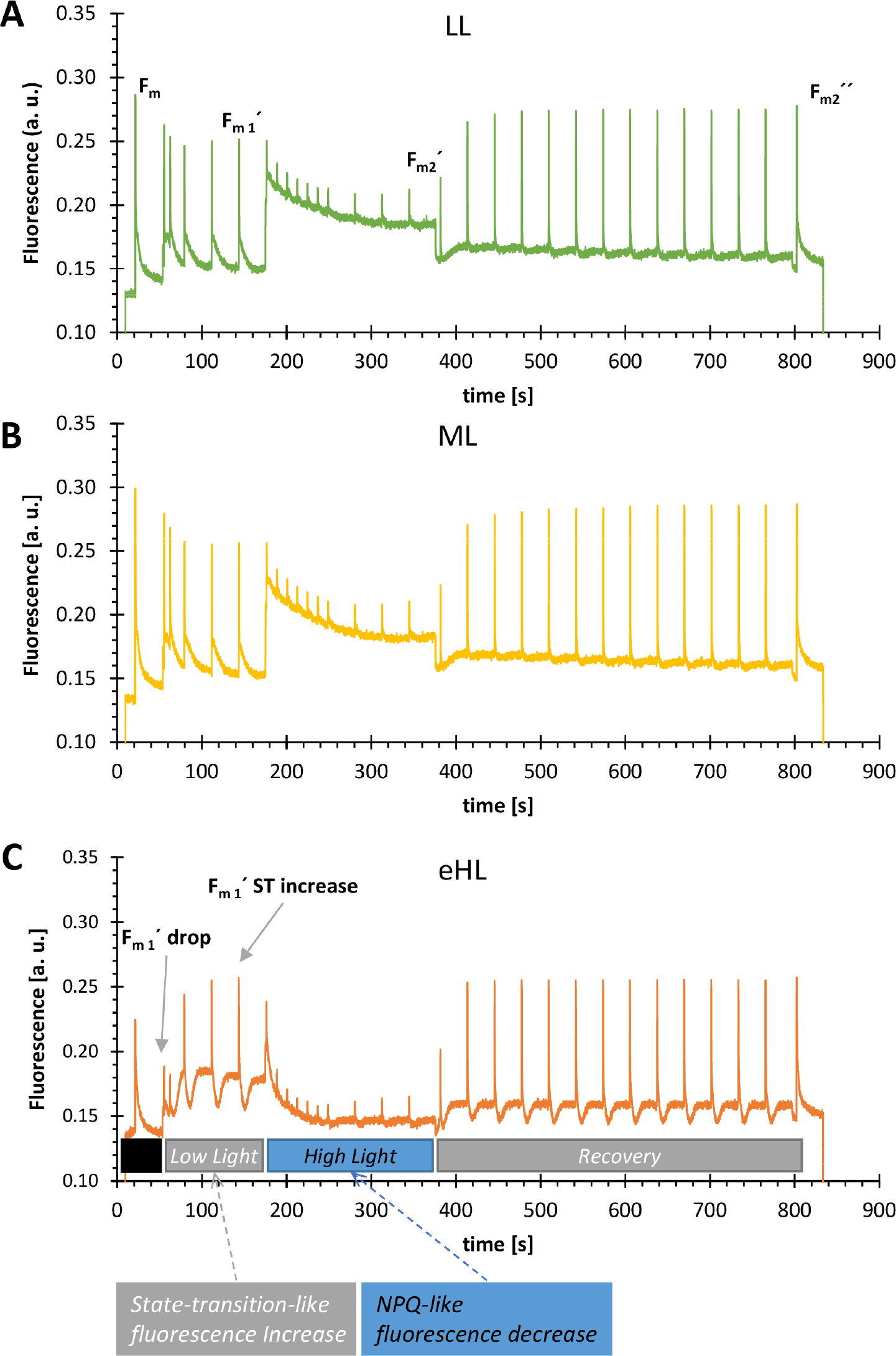
Fluorescence kinetic measurement under cellular acclimation to various light intensities. Fluorescence induction kinetics of dark-adapted cells were measured during a sequence of lights of different intensities. It included a dark period (black bar); low blue light period (81 μE, “Low light” bar); high blue light period (1374 μE, see “High light” bar); recovery period with low blue light (81 μE, “recovery” bar). The change in the maximal fluorescence (reflecting NPQ) was the measured regularly during multiple turnover flashes (4000 μE PAR) at the beginning of the protocol (dark-adapted F_m_ value), and during low light (F_m1_’) high-light (F_m1_’) and after recovery (F_m2_”). **A.** LL-adapted cells; **B.** ML-adapted cells; **C.** eHL-adapted cells; the sudden maximal fluorescence drop (“F_m1_’ drop”) and subsequent increase (“F_m1_’ state transition (ST) increase”) are indicated in panel C. Data represent the typical kinetic curves.

In eHL-adapted cells, only a transient decrease of F_m_ was observed in a few seconds of low-blue light illumination ( see “F_m1_’ drop” in Figure 5C) that was later overcome by the increase (see “F_m1_’ ST increase”), which is reminiscent of the State 1-to-State 2 transition previously observed in cyanobacteria (Calzadilla and Kirilovsky, 2020). The second type of NPQ induced by high-intensity blue light was present in all the cell samples analysed (LL, ML and eHL); however, the yield of this rapid quenching induced by high intensity blue light (see the second “blue” period in Figure 5) was markedly higher and exhibited the faster kinetics in eHL-adapted cells compared to LL and ML counterparts (see ‘high-light’ phase in Figure 5).

Overall, the *in vivo* dissection of the NPQ kinetics showed different strategies of photoprotection in LL- and ML-adapted cells in comparison to eHL cells; only the LL- and ML-adapted acido-thermophilic red algae seem to possess the previously described flash-induced NPQ detected in mesophilic red algae (Delphin et al., 1998) – a process that was absent in eHL-adapted cells.

### Photosynthetic microdomain formation during long-term varying light conditions

We studied the putative changes in the photosynthetic microdomain organisation inside thylakoids during long-term adaptation to varying light intensities (Figure 6). The microdomains represent thylakoid membrane regions with different compositions of pigment-protein complexes (photosystems and PBS) that were recently identified in cyanobacteria (Strašková et al., 2019). We conducted confocal fluorescence imaging of intact cells adapted to LL, ML, and eHL conditions (35, 90 and 350 μE PAR, respectively). In the previous works (Steinbach et al., 2015; Konert et al., 2019; Strašková et al., 2019; Canonico et al., 2020; Canonico et al., 2021), two main types of microdomains were identified in cyanobacteria – those resembling higher plant grana (predominantly with PSII complexes and associated antenna, i.e. PBS in the case of cyanobacteria) and those resembling plant stromal lamellae (more PSI and fewer PSII complexes) (see Strašková et al., 2019 for details). Therefore, we hypothesised that a similar variability in PBS and PSI spatial organisation occurs also in *C. merolae* thylakoids. To verify our hypothesis, we conducted confocal imaging of PSII + PBS (grana-like) microdomains and PSI (stroma-like) areas by detection of red-shifted fluorescence emission of PSI (see Materials and Methods). As shown in Figure 6A, PBS fluorescence was generally highly co-localised with the fluorescence of PSI (stromal lamellae-like areas); there was one exception when PSI was more abundant in the membrane areas with rather small PBS fluorescence (see green areas in Figure 6A) on the edges of the highly fluorescent PBS region (see yellow areas in Figure 6A). We define these high fluorescence regions as Microdomains Type I (MD-1, see Figure 6A) which are clearly visible in the heatmap representation of the PBS signal as white areas (see MD-1, Figure 6A). The remaining, rather smaller and less abundant areas contained more red-shifted emission signal originating from PSI (see MD-1, Figure 6A) which was caused by more homogenous distribution of red-shifted Chl*a* signal.

**Figure 6.**
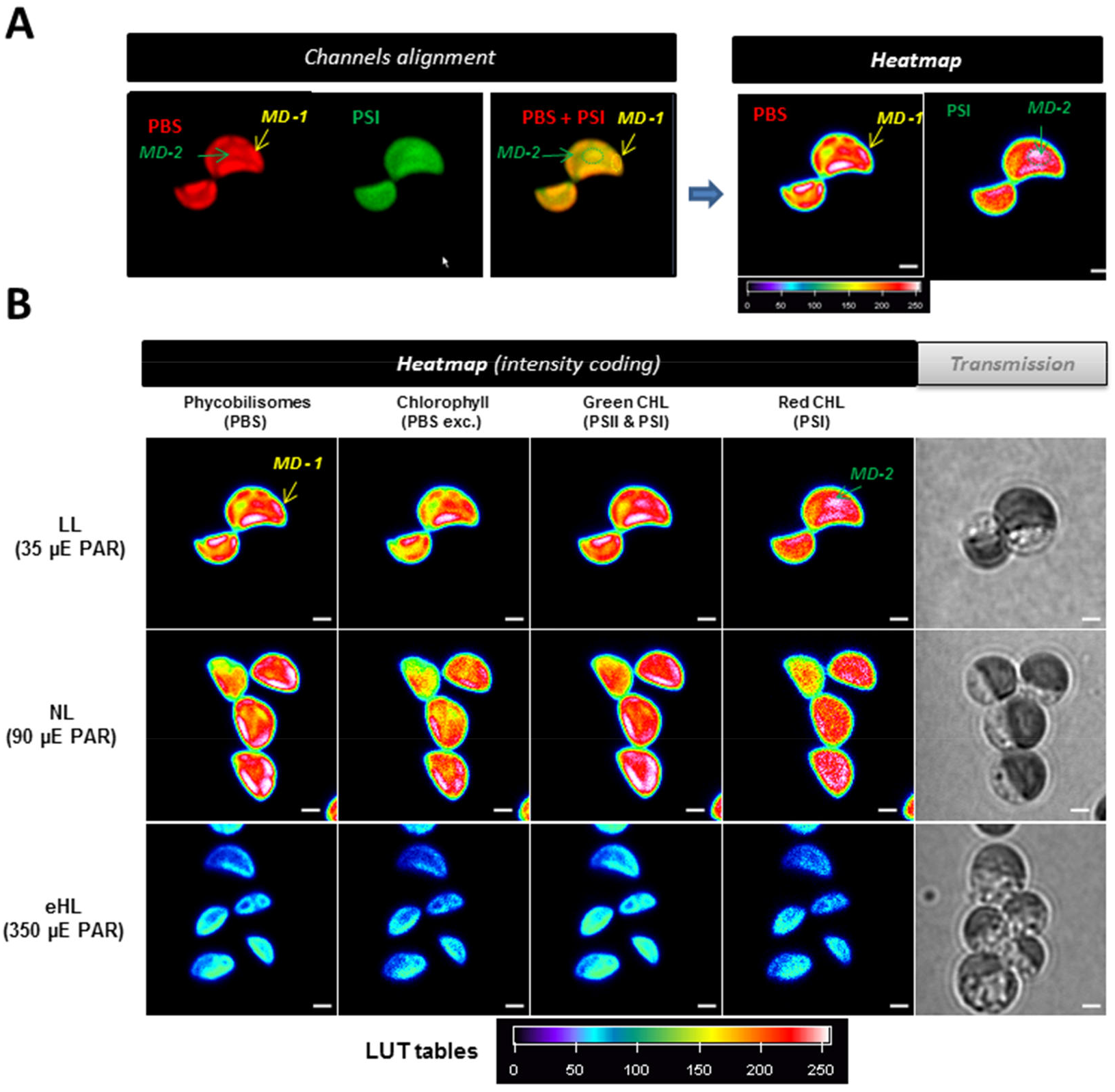
Light-dependent photosynthetic microdomain localisation in the *C. merolae* chloroplasts. Confocal fluorescence microscopy visualisation of chloroplasts was performed in cells subjected to 3-5-day light treatment of varying white light intensity (35, 90, 350 μE PAR, corresponding to LL, ML and eHL conditions). **Panel A.** Typical example of channels alignment (left part) and intensity coded heat map (right part) for two channels colocalisation: phycobilisomes (PBS; excitation at 594 nm, emission at 597-668 nm) and red-shifted Chl*a* emission (excitation at 488 nm, detection at 720-758 nm) reflecting red shifted PSI chlorophylls. PBS (in red) and PSI channels (in green) and their overlap to visualised microdomains (MD-1 or MD-2). Heat maps of PBS and PSI (right part) show heterogeneous intensities inside chloroplasts (from low intensity blue to high fluorescence white). **Panel B.** The heat maps of cells from LL, ML and eHL conditions and accompanied transmission picture. The intensity localisation of 4 channels were detected: (i)Phycobilisomes (PBS) (excitation at 594 nm, emission at 597-668 nm); (ii) Chl*a* emission (detected at 690-750 nm) upon PBS excitation wavelength (594 nm); (iii) Chl*a* emission from PSII (excitation at 488 nm, emission detection at 677-695 nm); (*iv*) red-shifted Chl*a* emission from PSI (excitation at 488 nm, detection at 720-758 nm). *Right*, corresponding light microscopy images of the cells. Scale bar: 1 μm. All the images of cells adapted to different illumination conditions (LL, ML and eHL) were acquired with the same microsocopy setup.

In the next step, we studied these structures in different light regimes. In all cases, we observed the asymmetric distribution of the fluorescence patterns for PBS and PSII that confirmed the presence of microdomains at LL, ML and eHL conditions; the microdomains of Type 1 (MD-1, see yellow in Figure 6) were the most abundant, whereas the MD-2 areas (with higher PSI occurrence) were localized in the edge of the MD-1 areas where the PSII and PBS emission was strongly decreased.

The MD-1 type (with more PBS and PSII signal) were visible at all light regimes, and they were more pronounced in the thylakoid membrane regions close to the edge of the chloroplast (see heatmap in Figure 6B). On the contrary, the red-shifted Chl*a* fluorescence (720-758 nm, a functional marker of the PSI complex) was more uniformly distributed throughout the chloroplast only under LL and ML conditions (Figure 6B). During eHL adaptation, MD-1 microdomains were predominantly formed, as judged by the co-localisation of fluorescence of all three photosynthetic complexes (PBS, PSII and PSI). Simultaneously, in eHL conditions the MD-1 areas were almost exclusively present at the edge of the chloroplast. It indicates remodelling of the photosynthetic microdomains under eHL stress, accompanied by the 2-fold reduction of chloroplast size compared to LL and ML grown cells.

In summary, we identified heterogeneous thylakoid membrane areas in *C. merolae* thylakoids containing predominantly the PSII-PBS MD-1 microdomains (functionally grana-like), in line with the study of Straskova et al. (2019). The areas with more dominant red-shifted Chl*a* emission signal (PSI areas corresponding to MD-2-type microdomains) bordered the MD1 microdomains, as the PSI fluorescence signal was more homogeneously distributed inside the thylakoids (Figure 6).

### Transcriptome changes in response to light quality and quantity adaptation in *C. merolae* cells

We hypothesised that the prolonged increased excitonic pressure on PSI (FRL treatment) or PSII (RL/eHL treatment) complexes may affect not only the relative antenna size (as observed for 77K fluorescence data, Figures 2 and 3) and stoichiometry of photosystems but it may also alter transcriptomic regulation of the genes encoding PSI/PSII subunits and related regulatory components involved in PSII assembly and dimerization, OEC formation, and the components of the antioxidant protective pathways. In order to verify these hypotheses, we performed a global transcriptome analysis by generating RNAseq libraries from total RNA isolated from cells adapted to varying light intensity and quality. We then dissected the statistically significant changes in the transcript levels of genes associated with these functions (see Table S1) using the ML transcriptome as the control for the eHL condition, and ML at time 0 as the control for cells at 48 h of RL and FRL adaptation. The results of the transcriptomic analysis are presented in Figure 7.

**Figure 7.**
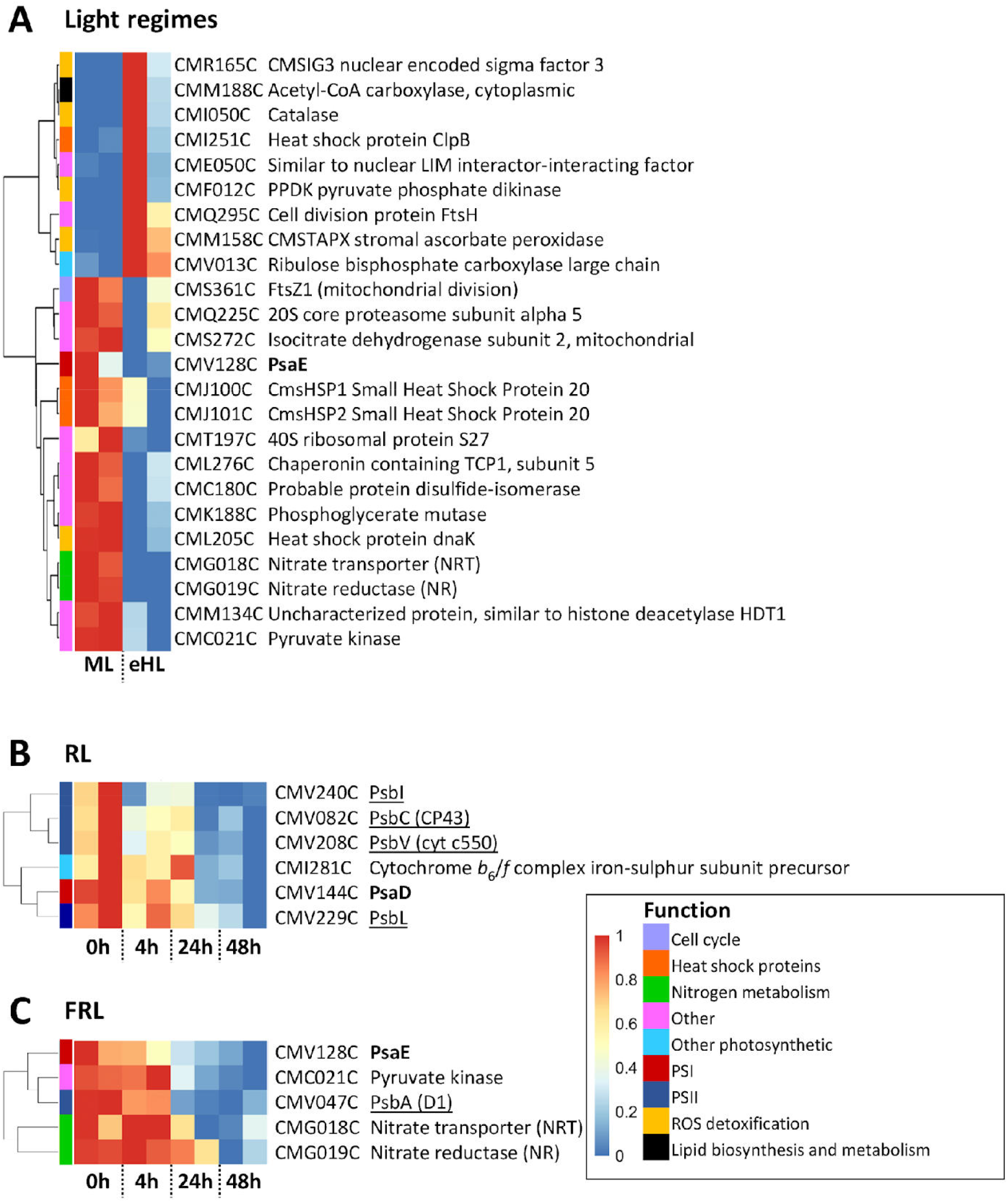
Light-dependent functional gene regulation analysis in *C. merolae*. **(A)** Comparison of transcriptome levels between the ML- and eHL-grown cells, shown are differential genes associated with indicated functions. *Top of the panel*: genes upregulated in eHL, compared to ML control. *Bottom of the panel*: genes downregulated in eHL compared to ML control. **(B)** Transcriptomic changes after RL treatment. Most significant changes occur after 24 h. All functionally associated genes that underwent significant expression level changes are directly related to photosynthesis. **(C)** Cultures undergoing FRL treatment. Each matrix represents two biological replicas (n=2). Maximum and minimum expression levels were used to normalise the data. Only statistically significant transcript level changes are displayed from the total set of transcripts, as shown by the clustering dendrograms on the left. *Blue*: transcript downregulation; *Red*: transcript upregulation (color scale shown in legend). *Inset*: color coding for the gene product function. On the right of each panel, similarity between each expression pattern is displayed. *Bold*: genes coding PSI subunits; *underlined*: PSII subunits; *italics*: genes encoding other components of electron transport pathways. Values in the heatmaps are min-max normalized (see Materials and Methods section).

#### eHL adaptation

We observed upregulation of the ROS detoxification genes (catalase, ascorbate peroxidase), transcripts encoding CO2 fixation enzyme (RuBisCO) and biosynthesis of fatty acids (ACC). Another upregulated gene encodes a key component of cell division (FtsH). Concomitant with these transcriptomic changes, we observed downregulation of the distinct small core subunit of PSI (PsaE). Significant downregulation of transcript levels was also observed for the mitochondrial division (FtsZ1), glycolysis (pyruvate kinase), and nitrogen metabolism enzymes (NRT, NR) (see Figure 7A).

#### RL adaptation

As expected, downregulation of the PSII core gene transcripts and the OEC components was detected, including CP43 inner antenna subunit of PSII, PsbI subunit involved in the assembly of CP43 with D1 reaction centre protein, PsbL associated with PSII dimer formation, and the OEC subunit PsbV (Figure 7B). Concomitantly, the transcripts encoding the PsaD subunit forming the ferredoxin binding pocket on the reducing side of PSI and cyt *b*6*f* Rieske subunit precursor were also downregulated.

#### FRL adaptation

Similarly to the RL-triggered transcriptomic downregulation of the PSI and PSII core subunits, FRL adaptation led to downregulation of PSI core subunit, PsaE (involved in ferredoxin binding) and PSII reaction centre subunit, D1 (Figure 7C). Importantly, we observed transcriptional downregulation of the downstream components of the central carbon and nitrate metabolism components, including NRT and NR enzymes involved in nitrate transport and nitrate reduction to nitrite as well as pyruvate kinase which a key enzyme of the glycolysis pathway (see Figure 7C).

## Discussion

Oxygenic phototrophs have evolved a remarkable plethora of strategies to react to changes in light intensity and spectral range, which allows them to thrive in a wide range of environmental conditions, from sun-soaked deserts to the deep shade of the rainforest and various depths in aquatic ecological niches. Varying light quality and quantity influences the balance between solar energy capture and utilisation in photosynthesis, affecting concomitantly the downstream processes of central carbon and nitrogen metabolism as well as cellular growth and division. High light stress often results in metabolic flux imbalances, photo-oxidative stress and/or slower development of plants or inhibition of microalgal cell division (Foyer and Noctor, 2005; Li et al., 2009b; Miyagishima and Tanaka, 2021; Sanchez-Tarre and Kiparissides, 2021).

In this work, we have performed comprehensive analysis of the extremophilic (acido-thermophilic) microalga *C. merolae*, using a combined and complementary spectroscopic, whole cell confocal fluorescence imaging, and global transcriptomic investigation in order to dissect the molecular mechanisms underlying the long-term adaptation mechanisms to varying light quantity and quality.

### Remodelling of the effective PSI and PSII antennae in C. merolae cells

This work has shown that significant remodelling of the effective antenna of PSII and PSI occurs in *C. merolae* cells adapted to varying light quantity and quality. Our absorption and 77K fluorescence analysis highlighted the significant remodelling of the antenna size of PSII and PSI during prolonged adaptation of *C. merolae* cells to LL, eHL, RL and FRL conditions (Figures 1–3). Several studies have confirmed increased accumulation of PBS in cyanobacteria and red algae in LL (PSII limited) conditions so that efficient photochemistry of PSII and PSI could be sustained (Kirst et al., 2014). Our current spectroscopic analysis confirms the higher accumulation of PBS proteins during LL adaptation compared to ML conditions; however, this trend was not sustained in RL and FRL conditions. This may be due to the fact that the actual PSII/PSI stoichiometry may favour PSII accumulation under RL and FRL conditions in *C. merolae* cells (similarly to eHL conditions) as a means to sustain high maximum quantum yield of PSII and balanced electron transport between PSII and PSI under light limiting conditions (see Figure 4).

Recent investigation of the *C. merolae* PSI-LHCI structure by X-ray crystallography (Antoshvili et al., 2019) and cryo-EM (Pi et al., 2018) provided an insight into the possible energy transfer pathways between the reaction centre complex and the associated LHCI antenna. In these structures, the core complex is associated with 3 or 5 Lhcr subunits forming 2 domains on the opposite side of the RC in the case of the larger PSI-LHCI supercomplex identified by cryo-EM (Pi et al., 2018). The three Lhcr subunits bind the core complex on the PsaF/PsaK side (Pi et al., 2018; Antoshvili et al., 2019), whereas the additional two Lhcr subunits identified in the cryo-EM structure of the *C. merolae* PSI-LHCI complex associate with the RC on the opposite side near the PsaI and PsaM core subunits (Pi et al., 2018). The specific pigment molecules identified in all the three structures were proposed to form the discrete energy transfer pathways. Among the proposed pathways, the most efficient ones most likely originate from the Chls present within Lhcr3 (denoted Lhcr3*) and Lhcr2 subunits in the larger PSI-LHCI cryo-EM structure, as these two Lhcrs not only have a close distance to the core (especially the distinct Chls in the PsaA subunit adjacent to Lhcr3) but they also converge with several other candidate pathways identified in the same structures, thus improving the energy transfer efficiency (Pi et al., 2018). The structural investigation of the putative energy transfer pathways in *C. merolae* PSI-LHCI isoforms highlighted the fact that these pathways differ significantly in the red algal complex compared to the higher plant counterpart in terms of the specific locations and pigments involved. It is therefore tempting to speculate that the distinctive energy transfer pathways may be generated upon remodelling of the pigment content of the peripheral LHCI antenna and associated reaction centre, as observed by red Chls accumulation in several recent studies (Haniewicz et al., 2018; Pi et al., 2018; Abram et al., 2020; Chang et al., 2020).

Indeed, we have recently shown by time-resolved fluorescence spectroscopy that the kinetics of energy transfer processes differ between *C. merolae* PSI-LHCI complexes isolated from LL, ML and eHL conditions. We showed that the average rate of fluorescence decay is not correlated with the size of LHCI antenna and is twice as fast in complexes isolated from ML-adapted cells (~25–30 ps) than from both LL and eHL cells (~50–55 ps) (Abram et al., 2020). We proposed that this difference is mainly due to a contribution of a long ~100-ps decay component detected only for the LL and eHL PSI-LHCI samples. We proposed that the lack of this phase in ML PSI-LHCI complexes is caused by the perfect coupling of ML LHCI antenna to the reaction centre and lack of low-energy Chls in this type of LHCI. On the other hand, the presence of the slow, ~100-ps, fluorescence decay component detected in the same work in LL and eHL PSI-LHCI complexes may be due to rather weak coupling between the respective PSI cores and LHCI antenna complexes, and due to the presence of particularly low-energy red Chls in LHCI.

Our studies on *C. merolae* cells and isolated PSI-LHCI complexes have revealed the remarkable functional flexibility of light harvesting strategies that have evolved in LL and eHL PSI-LHCI complexes in response to harsh or limiting light conditions involving accumulation of low energy Chls of a dual function: as energy traps or as FRL absorbing antenna pigments, respectively (Abram et al., 2020; present work). A similar observation was reported very recently for the HL PSI-LHCI complex in which decoupling of the LHCI antenna and reaction centre was detected by transient fluorescence decay spectroscopy (Chang et al., 2020).

### Strategies for fluctuating light utilisation in C. merolae cells

In this study, we have conducted a comprehensive analysis of the photosynthetic performance of *C. merolae* cells under fluctuating light. To this end, we characterised the three main parameters related to the efficiency of light utilisation by PSII for photochemistry and non-photochemical protective response. The first parameter, F_v_/F_m_ ratio, describes the maximum quantum efficiency of PSII photochemistry, which reflects the maximum efficiency of the PSII complex. The second parameter, ΦPSII, corresponds to actual PSII efficiency, and measures the proportion of absorbed photons used for photochemistry (Maxwell and Johnson, 2000). The third parameter, NPQ, characterizes the photoprotective mechanism of the non-photochemical quenching measured from the quenching of the maximal fluorescence in light. In a physiological context, the NPQ parameter monitors the apparent rate constant for heat loss from PSII (Baker, 2008).

The F_v_/F_m_ values remained constant and corresponded with those previously reported for *C. merole* and other rhodophytes (Krupnik et al. 2013). Surprisingly, ΦPSII values were quite low, with the lowest one observed at LL (Figure 4). The protocol used for measuring fluorescence included a 20 min period of dark adaptation of the cells, which may not be enough to eliminate state transitions. In cyanobacteria, algae and green plants State 1 is associated with the higher fluorescence yield, quantum efficiency of PSII and lower quantum efficiency of PSI. Conversely, in State 2 lower fluorescence yield and PSII quantum efficiency is observed, while PSI is characterized by its higher quantum efficiency (Minagawa, 2011). Lower F_v_/F_m_ and ΦPSII would suggest that in LL *C. merole* cells were in State 2. During this state excitation energy is redistributed to PSI (Campbell et al., 1998). Such assumption seems to be in congruence with the results of the absorption spectra analysis confirming bigger PSI antennae size at LL. Under moderate light, F_v_/F_m_ and ΦPSII values slightly, albeit significantly, increased, suggesting a necessity for balancing excitation energy, likely through gradual transition to State 1. This conclusion can be supported by the observed increase in the PSII/PSI Chl*a* emission ratio and/or increased energy transfer from PBS to PSII, as inferred from 77K fluorescence analysis. Moreover, the low ΦPSII values especially for LL cells could be also affected by the flash-induced quenching of maximal fluorescence that was observed in our study only in LL and ML cells (see Figure 5).

Mullineaux and Emlyn-Jones (2005) argued that ST process seems to be physiologically relevant only at LL intensities to ensure efficient utilization of absorbed light and plays no role as a photoprotective mechanism. In contrast, our present NPQ data supports the role of cyanobacterial-type ST in the initial phase of NPQ in eHL-adapted red algal cells, with the absence of this transient process in LL and ML cells (Figure 5). Tentatively, it could be proposed that under LL conditions *C. merole* cells prioritize light utilization of PSI, while not losing the photosynthetic capacity under eHL stress, as confirmed in our study by the observation of much higher ΦPSII values in eHL cells compared to LL counterparts.

The observation in eHL-adapted cells of transient fluorescence increase in low blue light and stronger fluorescence quenching upon the high blue light excitation (Figure 5) suggests the existence of two distinct mechanisms of NPQ in red algae. In eHL-adapted cells, the classical photoprotective response detected in continual high-blue light was the most dominant photoadaptive process in comparison to LL- and ML-adapted cells (see decrease in F_m2_’ value). On the contrary, the multi-turnover flash-induced NPQ, evoked with low-blue light in the background (see decrease of F_m_ to F_m1_’ in Figure 5), was only transiently observed in eHL-adapted cells (see F_m1_’drop in Figure 5C), whereas in LL- and ML-adapted cells this phenomenon was dominant. In eHL-adapted cells, the flash-induced decrease of the F_m_ parameter was compensated by state transitions detected by an increase of F_m_’ in low blue light (compare Fig. 5A, B and 5C).

The ST-related component of PSII fluorescence changes, observed in our study for eHL cells, is reminiscent ST in cyanobacteria, where illumination of dark adapted cells with low intensity blue light (mainly absorbed by Chl*a*, light 1) leads to the similar fluorescence increase (Canonico et al., 2021). It has already been proven in cyanobacteria that this phenomenon (i.e., the fluorescence increases upon low-intensity blue light illumination) is caused by the induction of low fluorescence State 2 in the dark (Papageorgiou et al., 2007). The subsequent fluorescence increase in low blue light is therefore connected with state transitions (Kana et al., 2012) due to the preferential excitation of PSI by low blue light leading to oxidisation of PQ pool (State 2-to-State 1 transition) (Calzadilla and Kirilovsky, 2020). These observations imply that in the dark, eHL cells were adapted to low-fluorescence State 2 (as it is typical for cyanabacteria), whereas LL- and ML-adapted cell were intrinsically in the high-fluorescence State 1 (as it is typical for higher plants and green algae) (see Kaňa et al., 2012; Papageorgiou et al., 2007). In cyanobacteria, the adaptation to these two photoprotective states has been rationalized by a difference in the redox state of the thylakoid membrane components due to the dark reduction processes driven by reductases (Kaňa et al., 2012; Papageorgiou et al., 2007). In addition, the PQ pool reduction can also occur due the activity of respiratory complexes like NAD(P)H dehydrogenase or/and succinate dehydrogenase that are localized in the thylakoid membrane of cyanobacteria (see e.g. Liu et al., 2012; Liu, 2016). It is not clear if a similar redox-state induced phenomenon occurs also in red algae. If it is the case it is likely to be connected with one of the chloroplast NAD(P)H dehydrogenases (Ma et al., 2021). Experiments are underway to address this issue.

Our data clearly prove that the molecular mechanism of fast photoprotective response in this extremophilic red algae is strongly dependent on the light intensity to which the cells are adapted in the long term. We detected dominant flash-induced NPQ mostly in LL- and ML-adapted cells. In eHL cells, the effect is only transient (see F_m1_’drop in Figure 5C) as the fluorescence quenching is compensated by ST-induced increase in fluorescence at low blue light (see “F_m1_’ ST increase” in Figure 5C). These data imply that the ST mechanism or regulation of excitation redistribution between photosystems is functionally more important in eHL-adapted cells. The mechanism of ST in red algae is still rather enigmatic, as it is often hard to resolve ST from the NPQ response (Delphin et al., 1996; Delphin et al., 1998). Nevertheless, it seems that ST in red algae most likely involve modulation of functional association (energetic coupling) of PBS and PSII (Ueno et al., 2015) or indirect energy transfer (spill-over) between PSII and PSI via PBS (Yokono et al., 2011; Ueno et al., 2017).

The LL- and ML-adapted red algal cells exhibit a typical NPQ response, i.e., PSII fluorescence quenching in high-intensity blue light, and no involvement of the ST induced fluorescence increase upon low blue light illumination. The typical NPQ response was observed through flash-induced quenching previously described for mesophilic red algae (Delphin et al., 1996; Delphin et al., 1998). The NPQ photoprotective response was previously shown to be triggered by low pH and to occur inside the PSII reaction centre acting as the quenching locus (Krupnik et al., 2013). In our study, this type of pH-dependent quenching was observed for all cell types (LL, ML and eHL) during illumination with high blue light (see decrease in F_m1_’ to F_m2_’ in Figure 5). The molecular mechanism of the second type of NPQ identified in our study, the MT flash-induced quenching (Delphin et al., 1998) does not require continuous high-light irradiation as it is present even at low blue light (see decrease from F_m_ to F_m1_’ in Figure 5). Its origin and molecular mechanism is more enigmatic, it requires relatively short pulses of high light (less than 1 second) and it is triggered by low pH and recovered by ATP synthase activity under conditions favouring pseudolinear and cyclic electron transport (see Delphin et al., 1998). Based on our data, it is tempting to speculate that this process could relate to the spatial redistribution of PSI observed in eHL and LL thylakoids (see Figure 6B). Subsequently, PSI may play a role in the rapid phase of the quenching (see Figure 5C), likely via excitation energy spill-over from overexcited PSII onto PSI that we postulate may be primed for quenching. This hypothesis is supported by the much faster quenching kinetics in eHL cells compared to LL and ML counterparts (see Figure 5).

### Formation and remodelling of the photosynthetic microdomains in C. merolae thylakoids

We have identified heterogeneous organisation of pigment-protein complexes (PPCs) (Strašková et al., 2019) into photosynthetic microdomains in red algae (Figure 6). These microdomain areas (MDs) were originally defined for cyanobacteria as specific thylakoid membrane regions with a diameter between 0.5-1 μm and characteristic ratios of the three PPCs, namely PSI, PSII and PBS (Strašková et al., 2019). The MDs in cyanobacteria and red algae correspond functionally to the structural thylakoid domains of higher plants and green algae, i.e. grana stacks and stromal lamellae, although no such structures exist in red algal chloroplasts. The common feature of MDs and appressed/non-appressed regions of thylakoids is the heterogeneity of PPC distribution, with more PSII and its associated antenna present in the MD-1 microdomains and appressed lamellae, whereas PSI with its associated antennae present in the MD-2 (red algae) and non-appressed regions (green algae and higher plants). As we did not used fluorescence tagging of PSI, as it was applied for cyanobacterial MDs (Strašková et al., 2019), we identified PSI localization based on red-shifted signal originating from PSI emission at room temperature (in the range 720-758 nm, Figure 6) and also at low temperature (see 77K emission spectra in Figures 2 and 3). Previously we used a similar approach for cryo-microscopy of cyanobacterial (*Anabaena* sp. PCC 7120) cells (Steinbach et al., 2015) which served as a proof-of-concept for the present study. Here, we have found that PSI seems to be more homogeneously distributed in *C. merolae* thylakoids. The heterogeneous MD distribution was clearly visible in thylakoids of *C. merolae* based on the PSII-PBS fluorescence signal (MD-1 domains marked in Figure 6). Conversely, PSI was more uniformly distributed throughout thylakoids compared to PSII-PBS, albeit more abundantly at the border of the MD-1 domains. These PSI-rich regions of thylakoids were defined as the MD-2 domains in our study (see in Figure 6A). This heterogeneity of PSI distribution was abolished in eHL thylakoids, when all PPCs were rather uniformly distributed (Figure 6B). We hypothesise that this phenomenon, observed exclusively in eHL conditions, is based on PSI intermixing with PSII-rich microdomains. Such redistribution of PSI in long-term eHL adaptation could alleviate excitonic pressure on PSII by: (1) triggering State 2-to-State 1 transition and (2) increasing the overall NPQ response and/or PBS-PSII-PSI energy spill-over (Ueno et al., 2017; Yokono et al., 2011). In support of this hypothesis, the ST component in low blue light and increased NPQ response in orange light were observed in our study in eHL-adapted cells (see Figure 5), as discussed above. In the previous work, MDs in cyanobacteria were found to be stable in the range of minutes (Straskova et al., 2019) and their distribution changes slowly under fluctuating light conditions on a timescale from hours to days (Canonico et al., 2020). This seems to be also the case for the red-algal MDs, as PPCs, including PBS, are almost immobile in the extremophilic microalgae (Kaňa et al., 2014).

In summary, our present data on redistribution of red algal MDs in fluctuating light open new avenues to study the role of subcellular heterogeneity in thylakoid membranes in maintaining the highest photosynthetic efficiency under light stress conditions.

### Transcriptomic feedback loop between light reactions and central carbon and nitrogen metabolism

In higher plants, acclimation to HL involves increases in electron transport and carbon metabolism, but no change in the abundance of photosynthetic reaction centers. (Miller et al., 2017). Indeed, our present data show increased PSII effective quantum yield with increased white light intensity and in RL conditions (Figure 4), highlighting the improved linear electron transport across PSII in the presence of sustained PSII overexcitation. Yet, at the global transcriptomic level, downregulation of the PSII and PSI subunits involved in charge separation and charge transfer (D1), OEC regulation (PsbV), and ferredoxin binding (PsaE and PsaD) occurs in *C. merolae* cells under the same illumination conditions (see Figure 7). This suggests the existence of a subtle interplay between transcriptomic regulation of the photosynthetic components and the modulation of their functionality.

In the present study, we show upregulation of transcripts closely related to regulation and protection of photosynthetic light reactions in eHL-adapted *C. merolae* cells. These include the components of the ROS detoxification pathways (catalase, ascorbate peroxidase), CO2 fixation (RuBisCO), and biosynthesis of fatty acids (acetyl-CoA carboxylase, ACC). Another eHL-upregulated gene encodes a postulated component of cell/mitochondrial division (FtsH; Itoh et al., 1999). Concomitant with these transcriptomic changes, we observed downregulation of the distinct small core subunits of PSI in line with previously published proteomic data showing a partial or complete loss of several core subunits of the eHL PSI-LHCI complex (PsaE, PsaK, Haniewicz et al., 2018). Significant downregulation of transcript levels was also observed for the mitochondrial division (FtsZ1), glycolysis (pyruvate kinase), and nitrogen metabolism enzymes (NRT, NR) (see Figure 7) pointing towards a close interplay between regulation of long-term adaptation of the photosynthetic apparatus to eHL and the downstream metabolic pathways of central carbon and nitrogen metabolism.

In line with our transcriptome regulation observations, a recent quantitative proteomic study on HL-acclimated *Arabidopsis thaliana* plants, which included a physiological study of photosynthetic performance parameters, has elucidated the upregulated components and processes involved in the acclimation process, including increased CO2 assimilation and PSII electron transfer rate (ETR (II)) values (Flannery et al., 2021). The observed increased linear electron transport correlated in turn with the increase in protein levels for ATP synthase, cyt *b*6*f*, FNR2, TIC62 (FNR thylakoid tethering protein) and PGR6 components, thereby augmenting the PQ pool and forming the cyclic electron flow (CEF) molecular machinery, resulting in increased ATP production (Flannery et al., 2021). The same study has shown that improved photosynthetic capacity in HL-grown plants is paralleled by increased CEF, which positively correlated with the abundance of NDH, PGRL1, FNR1, FNR2 and TIC62 (not PGR5) components of CEF. On the other hand, LL-grown plants favoured a slowly reversible NPQ component (qI reversible photoinhibition component in fluorescence quenching measurements), which positively correlated with LCNP abundance (component of NPQ). HL-grown plants favoured the rapidly reversible qE component of NPQ, which positively correlated with the abundance of a PsbS component involved in triggering the NPQ process. The long-term adjustment of the grana diameter positively correlated with LHCII levels, while grana stacking negatively correlated with CURT1 (thylakoid curvature protein) and RIQ protein (negative regulator of the grana size) levels.

Several other proteomic analyses in HL-adapted plants have shown that the total content of the photosynthesis-related components, i.e. RuBisCo and the cytochrome *f* component of the cytochrome *b*6*f* complex, increases upon exposure of plants to increased incident light (Yin and Johnson, 2000; Athanasiou, 2007; Athanasiou et al., 2010). A close interplay between photosynthesis and downstream metabolic components was demonstrated in long-term HL-treated *Arabidopsis* plants in line with the transcriptomic data of our study. Microarray analysis of such plants combined with mutagenesis analysis have shown the crucial importance of the glucose-6-phosphate/phosphate translocator GPT2 for efficient photosynthetic performance, pointing towards a signalling role of this downstream component postulated to affect the partitioning of sugar phosphates between the chloroplast stroma and the cytosol, or shifting the phosphate balance of the cell (Athanasiou et al., 2010).

During RL adaptation, we observed transcriptional downregulation of the PSII inner antenna subunit, CP43, PsbI and PsbL core subunits involved in the assembly of the functional PSII dimer, and the OEC component, PsbV (Figure 7). Concomitantly, the transcripts encoding the PsaD subunit forming the ferredoxin binding pocket on the reducing side of PSI and cyt *b*6*f* Rieske subunit were also downregulated, likely constituting the protection mechanism preventing light-induced over-reduction of PQ pool under the conditions of overexcitation of PSII. Interestingly, no long-term changes in transcript levels were observed for the subunits constituting the functional antennae of photosystems during RL illumination. This suggests that for the maintenance of the highest photosynthetic performance in *C. merolae* cells exposed to long-term RL illumination, no metabolic energy investment is allocated to *de novo* synthesis of light harvesting components. In contrast to *C. merolae*, long-term application of varying light quantity and quality of *Chlamydomonas reinhardtii* cells has led to upregulation of the RuBisCo small subunit under red phototrophic growth, implying an increased demand for energy generation in this green microalga required to establish a shorter cell cycle and maintain a rapid duplication rate in conjunction with increased photosynthetic efficiency as well as reduced mutual shading across the light path (Sanchez-Tarre and Kiparissides, 2021). The same study showed that metabolic energy investment is based on transcriptomic upregulation of fatty acid biosynthesis and cell cycle components, which ensures shortening of the cell cycle and maintenance of a rapid duplication rate (Sanchez-Tarre and Kiparissides, 2021). In our work, transcriptomic upregulation of fatty acid biosynthesis and cell cycle enzymes was observed for eHL adaptation rather than adaptation to light preferentially absorbed by PSII (RL), indicating the subtle regulatory differences between these two groups of evolutionary distant microalgae.

Surprisingly, the transcriptome of the FRL-adapted cells was characterised by downregulation of PSI core subunit, PsaE (involved in ferredoxin binding) and PSII reaction centre subunit, D1 (Figure 7), implying a negative feedback loop between transcriptional regulation of PSI and its effective quantum efficiency (compare Figures 4 and 7). Similarly to eHL adaptation, transcriptional downregulation of the downstream components of the central carbon and nitrate metabolism components was also observed for the FRL adapted cells, including NRT and NR enzymes involved in nitrate transport and nitrate reduction to nitrite as well as pyruvate kinase which a key enzyme of the glycolysis pathway (see Figure 7). The observed robust expression of NR and NRT genes under non-stress conditions (ML, early time points of RL and FRL treatment) and their downregulation under light limiting conditions (eHL and later time points of RL and FRL treatment) are in contrast with the previous report (Fujiwara et al., 2015) on the tight regulation of these genes in the standard growth medium based on ammonia, such as that used in this study (Fujiwara et al., 2015). The discrepancy between our results and the previous ones requires further study.

In summary, the present study has identified several important molecular mechanisms underlying the long-term adaptation of the *C. merolae* cells to varying light conditions. They include: (1) remodelling of the effective antenna size of both photosystems; (2) rearrangement of the PSB/PSII/PSI microdomains within thylakoids; (3) modulation of the photosynthetic performance parameters as the means for optimal light utilisation, especially by utilising two distinct NPQ mechanisms in eHL versus LL/ML conditions and (4) transcriptional regulation of photosynthesis, and its regulatory components and downstream metabolic pathways including ROS detoxification pathways, cell/organelle division, as well as carbon and nitrogen metabolism (Figure 8). Such an intricate network of interplay between light-driven reactions and downstream metabolic pathways provides the strong basis for maintaining the highest photosynthetic performance under fluctuating light. The efficient and dynamic adaptation of *C. merolae* cells to eHL conditions using multiple molecular pathways identified in our study is likely to be related to the natural habitat of this extremophilic microalga, i.e., humid shallow hot springs (Ciniglia et al., 2004), whereby high light stress is prevailing.

**Figure 8.**
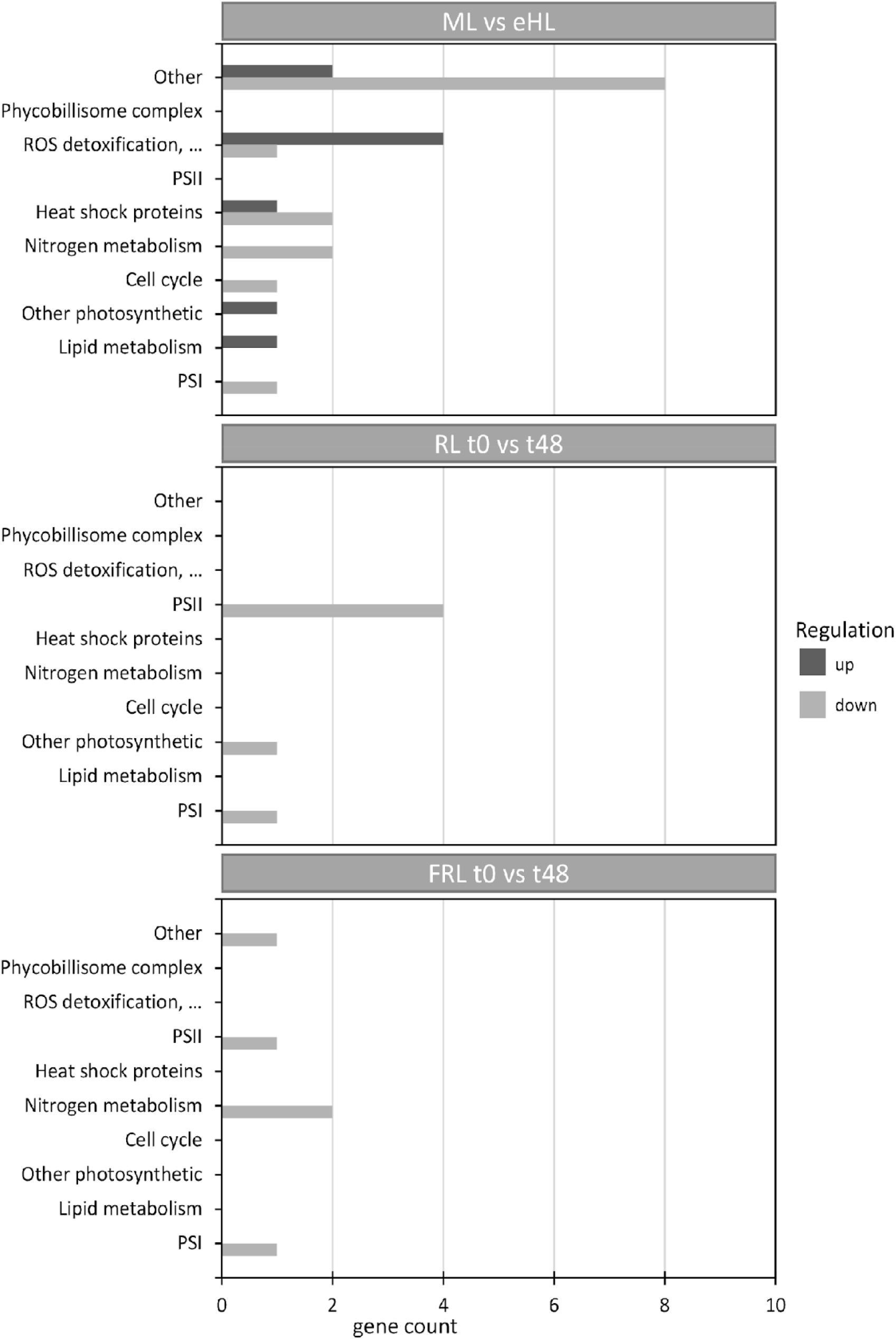
Overview of the metabolic processes affected by varying light conditions applied to the *C. merolae* cells derived from the total gene expression patterns. Varying light quantity and quality exerts a variety of adaptive responses in *C. merolae* cells related to regulation of photosynthesis, ROS detoxification pathways, lipid biosynthesis and metabolism, cell/organelle division, as well as central carbon and nitrogen metabolism.

## Materials and Methods

### Culture growth conditions and sample harvesting

*Cyanidioschyzon merolae* (strain NIES-1332, National Institute of the Environmental studies, Japan) cultures were grown as described in (Haniewicz et al., 2018) in Allen-2 medium (Minoda et al., 2004) at pH 2.5 and 42 °C, in constant white light with 5% CO2 bubbling. For varying white light intensity adaptation experiments, cells were grown in three different light regimes: LL (low light) of 35 μmoles photons m^−2^ s^−1^ (μE) photosynthetically active radiation (PAR); ML (moderate light) of 90 μE PAR; and eHL (extreme high light) of 350 μE PAR in the growth chamber supplemented with fluorescent lights (Panasonic). Cells grown in the eHL regime were initially grown in 90 μE PAR for ~26 h with a three-step increase in light intensity every 18 h from 90 μE to 350 μE PAR. The cell culture growth in the final illumination conditions was conducted for a further 30 h. For varying spectral quality adaptation experiments, cell suspensions were grown initially in the fluorescent white light chamber in 60 – 65 μE PAR, after which they were transferred to the LED-illuminated chamber (Biogenet) (Suppl. Fig. 1) and grown for a further 48 h using 60 – 68 μE PAR of the desired spectral range (see Supplementary Figure 1). For Red Light (RL) experiments the light setup was: 2% Cold White (6500 K), 1% Warm White (2700 K), 100% Red (630-650 nm), 100% Deep Red (650-670 nm). For the Far-Red Light (FRL) experiments, the illumination conditions were: 25% White (6500 K), 14% Warm White (2700 K), 100% FRL (710-740 nm) (see Supplementary Figure 1). For the transcriptomic analysis, samples were harvested in 50 ml volume, centrifuged at 100 rcf for 15 min at 4 °C, after which the supernatant was discarded, cell pellet was weighed, then snap-frozen in liquid nitrogen, and stored at −80 °C until further use.

### Room temperature absorption

Absorption measurements were carried out according to the procedure described in Haniewicz et al. (2018), using a UV-1800 Shimadzu spectrophotometer with the TCC-100 temperature-controlled cell holder, using a quartz cuvette with an optical path length of 10 mm. The minimum full width at half maximum (FWHM) parameter was 1 nm. The detector wavelength resolution was 0.5 nm. For each measurement, 1 ml cell suspension was collected from each culture. Averaged spectra (n=2) were normalised to the PSI/PSII Chl absorption peak.

### Chlorophyll *a* concentration measurement

Chl*a* concentration measurements were estimated based on (Porra et al., 1989) in an experimental procedure described in Haniewicz et al. (2018). Briefly, 1ml of the culture was centrifuged (14,400 rcf, RT, 5 min). 900 μl of the supernatant was removed and the cells were resuspended in 900 μl of 80% acetone and vortexed for 2 min. and then centrifuged (14,400 rcf, RT, 5 min). 650 μl of the supernatant were transferred to the quartz cuvette and the whole spectra were measured. Chl*a* concentration was calculated by dividing the Chl absorbance at 680 nm by the extinction coefficient ε= 86.3 μg^−1^ μl^−1^ cm^−1^ (Porra et al., 1989).

### PAM measurements

PAM fluorescence measurements were performed with the FMS1 PAM system (Hansatech, Norfolk, United Kingdom) equipped with an amber light (594 nm) emitting diode as a source of measuring light. An internal halogen lamp (8 V/20 W) provided actinic and saturating light. Measurements were carried out with a 5.5-mm-diameter fiberoptic kept perpendicularly to the sample at a distance of 26 mm. Cell suspensions (1 ml) were drop-casted on the filter membrane, then placed onto a cylindrical holder after which an additional drop (~0.2 ml) of the medium was added to prevent desiccation. Subsequently, the holder was placed inside the DW2/2 chamber (Hansatech, Norfolk, United Kingdom) attached to a circulating water bath in order to maintain constant temperature set to 30 °C. Prior to measurements, cells were dark-adapted for 20 min (Bailey et al., 2005). Next, the saturation pulse of >3000 μE m^−2^ s^−1^ was applied in order to measure minimum (F0) and maximum (F_m_) fluorescence. Subsequently, actinic light, of intensity matching the culturing light conditions, i.e., 35, 90 and 350 μE m^−2^ s^−1^, was switched on. After reaching a steady-state another saturation pulse was provided so as to measure steady-state (Fs) and maximum (F_m_’) fluorescence in the light-adapted state. Obtained values allowed calculating the maximum (F_v_/F_m_=[F_m_-F_o_]/F_m_) and effective (ΦPS2=[F_m_’-F_s_]/F_m_’) quantum efficiency of PSII, and non-photochemical quenching (NPQ=[F_m_-F_m_’]/F_m_’) (Genty et al. 1989, Schreiber et al. 1986). The p-values of the observed parameter changes were calculated in the range of 0.007-0.03.

### Measurement of NPQ kinetics

The fluorescence kinetics and non–photochemical quenching (NPQ) of maximal fluorescence (F_m_) was measured by DUAL-PAM 100 fluorometer (Heinz Walz GmbH, Effeltrich, Germany) by a Dual-DR detector head (excitation 620 nm, detection above 700 nm) emitting measuring red light (20 Hz, 7 μmol photons m^−2^ s^−1^, λ = 620 nm). Cells were dark adapted for 10 min prior to measurements, and kinetic changes in F_m_ were measured immediately after dark adaptation. The four values of maximal fluorescence were obtained: (1) maximal F_m_ value of a dark adapted sample; (2) F_m1_’ value after 120 s of low intensity blue light illumination (81 μmol photons m^−2^ s^−1^, λ = 460 nm) followed with intense red flashes illumination (4000 μmol photons m^−2^ s^−1^, 400 ms long, λ = 620 nm); (3) F_m2_’ value after 200 s of high intensity blue light (1374 μmol photons m^−2^ s^−1^, λ = 460 nm); (4) Recovery value F_m2_’’ after 420 s at low intensity blue light (81 μmol photons m^−2^ s^−1^, λ = 460 nm). The used protocol followed the standard procedure used for cyanobacteria, as described in Canonico et al. (2021). The maximal value of fluorescence was estimated by a multiple turnover flash (red light λ = 620 nm, 4000 μmol photons m^−2^ s^−1^, duration 400 ms) every 10–30 s of the protocol.

### Steady-state 77K fluorescence measurements

Low temperature (77K) fluorescence emission spectra were measured with an LS55 fluorimeter (Perkin Elmer, Waltham, MA, USA) as described in Haniewicz et al. (2018). The cell suspension (3 μg Chl) was immediately mixed with 80% glycerol, then transferred to a quartz cuvette and inserted into the holder immersed in liquid nitrogen. Fluorescence was recorded after 4-5 min with horizontal and vertical polarisation filters in the optical path and the wavelength resolution of 0.5 nm. Chl*a* fluorescence was measured by excitation at 435 nm in the range of 450-800 nm with 515 nm cut-off filters in the optical path between the sample and detector. Excitation spectra were recorded by the fluorescence detection at 728 nm (PSI emission) with the excitation wavelength range of 400-700 nm. For all the measurements, light slits were adjusted for the optimal peak fluorescence signal value (500-800 a.u.). Emission spectra were normalised at 685 nm, while the excitation spectra were normalised to the PSI emission peak at 670 – 680 nm. All the spectra were obtained from 2 independent biological samples (n=2).

### *In vivo* confocal fluorescence measurements

The confocal pictures were acquired with an inverted Zeiss LSM 880 laser scanning confocal microscope (Carl Zeiss Microscopy GmbH, Oberkochen, Germany) equipped with a Plan-Apochromat 63x/1.4 NA Oil DIC M27 immersion objective. Phycobilisomes emission was excited by a HeNe laser (594 nm) and detected in 597 – 668 nm range by GaAsP detector. The Chl*a* autofluorescence was excited either directly at 488 nm (Argon laser) or through energy transfer via PBS excitation at 594 nm (by HeNe). The Chl*a* fluorescence signal was detected in two setups: (1) Whole range detection of Chl*a* emission (emission of PSI and PSII) by PMT detector between 690 – 750 nm, the fluorescence was excited through PBS excitation at 594 nm (by HeNe); (2) Double intervals detection of Chl*a* emission in blue- (677-695 nm; GaAsP detector, gain 790) and red-shifted regions (720 – 758 nm; PMT detector, gain 850); with excitation at 488 nm (Argon laser). The pinhole was adjusted for 1 AU for all channels, images were acquired with 177 μs pixel dwell time (pixel x-y scaling 81 nm per 81 nm), in five channels (image x-y size 128×128, 8-bit resolution). The transmission pictures were acquired by the T-PMT detector for 488 nm excitation (Ar laser). Pictures were analysed in ZEN black software (Carl Zeiss Microscopy GmbH, Oberkochen, Germany) and in Image J software (Schneider et al., 2012).

### Transcriptomic analysis

#### Total RNA Isolation

Cell pellets were resuspended in 500 μL cold phenol RNA lysis buffer (0.5 M NaCl, 0.2 M TrisoHCl (pH 7.5), 0.01 M EDTA, 1% SDS) and were sonicated briefly to shear the genomic DNA. Total RNA was extracted two times with acid phenol/chloroform (pH 4.5, Ambion), followed by chloroform extraction. RNA was EtOH-precipitated and resuspended in dH2O and then treated with Turbo DNase (Ambion) following the manufacturer’s instructions.

#### Strand-specific messenger RNA-sequencing

Qualities of total RNA samples were determined using an Agilent Bioanalyzer RNA Nanochip and arrayed into a 96-well plate (Thermo Fisher Scientific). Polyadenylated (PolyA+) RNA was purified using the NEBNext Poly(A) mRNA Magnetic Isolation Module (E7490L, New England Biolabs) from 1000 ng total RNA. Messenger RNA selection was performed using NEBNext Oligod(T)25 beads (NEB) incubated at 65 °C for 5 min followed by snap-chilling at 4 °C to denature RNA and facilitate binding of poly(A) mRNA to the beads. mRNA was eluted from the beads in Tris-HCL buffer, pH 7 incubated at 80 °C for 2 min then held at 25 °C for 2 min. RNA binding buffer was added to allow the mRNA to rebind to the beads, mixed 10 times and incubated at room temperature for 5 min. The sample plate was placed on the magnet and the supernatant discarded. The mRNA bound beads were washed twice then cleared again on magnet. The supernatant was again discarded, and mRNA eluted from the beads in 20 μL Tris-HCl, pH 7 incubated at 80 °C for 2 min. Total mRNA was transferred to a new 96-well plate.

First-strand cDNA was synthesized from heat-denatured purified mRNA using a Maxima H Minus First Strand cDNA Synthesis kit (Thermo-Fisher, USA) and random hexamer primers at a concentration of 200 ng/μL along with a final concentration of 40 ng/μL actinomycin D, followed by PCR Clean DX (Aline Biosciences) bead purification on a Microlab NIMBUS robot (Hamilton Robotics, USA). The second strand cDNA was synthesized following the NEBNext Ultra Directional Second Strand cDNA Synthesis protocol (New England Biolabs) that incorporates dUTP in the dNTP mix, allowing the second strand to be digested using USERTM enzyme (NEB) in the post-adapter ligation reaction and thus achieving strand specificity.

The cDNA was fragmented by Covaris LE220 sonication to achieve 250-300 bp average fragment lengths. The paired-end sequencing library was prepared following the BC Cancer Genome Sciences Centre strand-specific, plate-based library construction protocol on a Microlab NIMBUS robot (Hamilton Robotics, USA). Briefly, the sheared cDNA was subject to end-repair and phosphorylation in a single reaction using an enzyme premix (New England Biolabs) containing T4 DNA polymerase, Klenow DNA Polymerase and T4 polynucleotide kinase, incubated at 20 °C for 30 min. Repaired cDNA was purified in 96-well format using PCR Clean DX beads (Aline Biosciences, USA), and 3’ A-tailed (adenylation) using Klenow fragment (3’ to 5’ exo minus) and incubation at 37 °C for 30 min prior to enzyme heat inactivation. Illumina PE adapters were ligated at 20 °C for 15 min. The adapter-ligated products were purified using PCR Clean DX beads, then digested with USERTM enzyme (1U/μL, NEB) at 37 °C for 15 min followed immediately by 10 cycles of indexed PCR using NEBNext Ultra II Q5 DNA Polymerase (New England Biolabs) and Illumina’s PE primer set. PCR parameters: 98 C for 1 min followed by 10 cycles of 98 °C, 15 s; 65 °C, 30 s; and 72 °C, 30 s, and then 72 °C for 5 min. The PCR products were purified and size-selected using a 1:1 PCR Clean DX beads-to-sample ratio (twice), and the eluted DNA quality was assessed with Caliper LabChip GX for DNA samples using the High Sensitivity Assay (PerkinElmer, Inc. USA) and quantified using a Quant-iT dsDNA High Sensitivity Assay Kit on a Qubit fluorometer (Invitrogen) prior to library pooling and size-corrected final molar concentration calculation for Illumina HiSeq2500 sequencing with paired-end 75 base reads.

#### Bioinformatic analysis

We used fastp version 0.20.1 (Chen et al., 2018) to control and improve the sequence read quality before starting downstream analysis. This entailed general quality profiling followed by removal of low-quality reads, trimming of low-quality bases and adapter removal. The processed fastq files obtained from fastp were used for the read alignment.

#### Read alignment

Alignment of the trimmed reads was performed using STAR version 2.7.5c (Dobin et al., 2013) with the quantMode GeneCounts argument for producing gene counts. Remaining arguments were set to default during both the STAR indexing and alignment steps. Reads were aligned to the reference genome obtained from the *Cyanidioschyzon merolae* Genome Project v3 (Nozaki et al., 2007) using a custom annotation based on the Ensembl gene annotation ASM9120v1. The aligned reads were then sorted and indexed using samtools version 1.12 (Li et al., 2009a) before performing differential gene expression analysis.

#### Differential gene expression analysis

Read counts produced by the STAR aligner were collected as a count matrix that was then used as input for differential gene expression analysis using DESeq2 version 1.28.1 (Love et al., 2014). DESeq2 was run with fitType = “local” but with remaining arguments set to default. Adaptive shrinkage (Stephens, 2017) was used for correcting the expression differences observed for lowly expressed genes. Significant differential genes were identified by filtering on adjusted p-value ≤ 0.01 and absolute log2-fold change ≥ 0.75.

#### Functional gene regulation analysis

Functional gene regulation was investigated by intersecting the set of significantly differential genes with a set of genes that had been previously annotated with the specific biological function (see Table 1, Supplementary Material). The results were then visualized by plotting the number of significant differential genes per comparison split by regulation (up/down) and grouped by biological function.

**Table 1.**
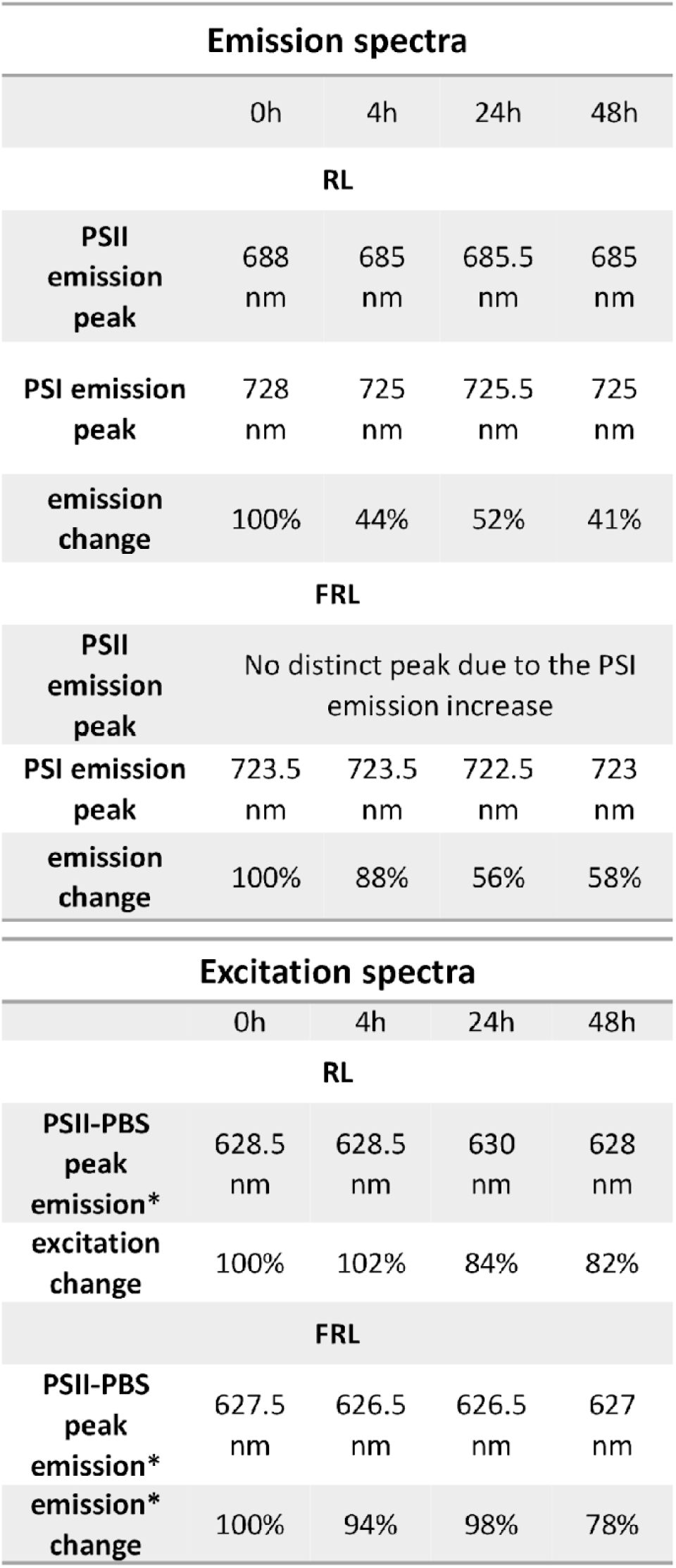
Emission peak wavelengths derived from the 77K emission and excitation spectra.

#### Visualization of gene expression

Gene expression results were visualised as heatmaps using the R package “Pheatmap” version 1.0.12 (https://github.com/raivokolde/pheatmap). DESeq2 normalized read counts were min-max normalized per gene across all compared conditions before being used for heatmap generation. Where applicable, genes were clustered by Euclidean distance.

DESeq2 normalized gene counts were visualized as boxplots for a subset of genes, selected on the basis of their differential expression upon each light treatment, by aggregating the counts from all replicates. The aggregated counts were independently plotted as boxplots for the comparisons *medium light versus extreme high light*, *medium light versus neutral pH* and the red light / far red-light time courses.

## Acknowledgments

JK and MA gratefully acknowledge support from the Polish National Science Centre (OPUS 8 grant no. UMO-2014/15/B/NZ1/00975 to JK). Part of this work was supported by the National Science Centre OPUS17 grant (UMO-2019/33/B/NZ3/01870 to JK). SR was supported by GenomeBC Sector Innovation Program grant RC18-3517 and NSERC Discovery Grant 298521.

## Supplementary Figures and Tables

**Supplementary Figure 1.**
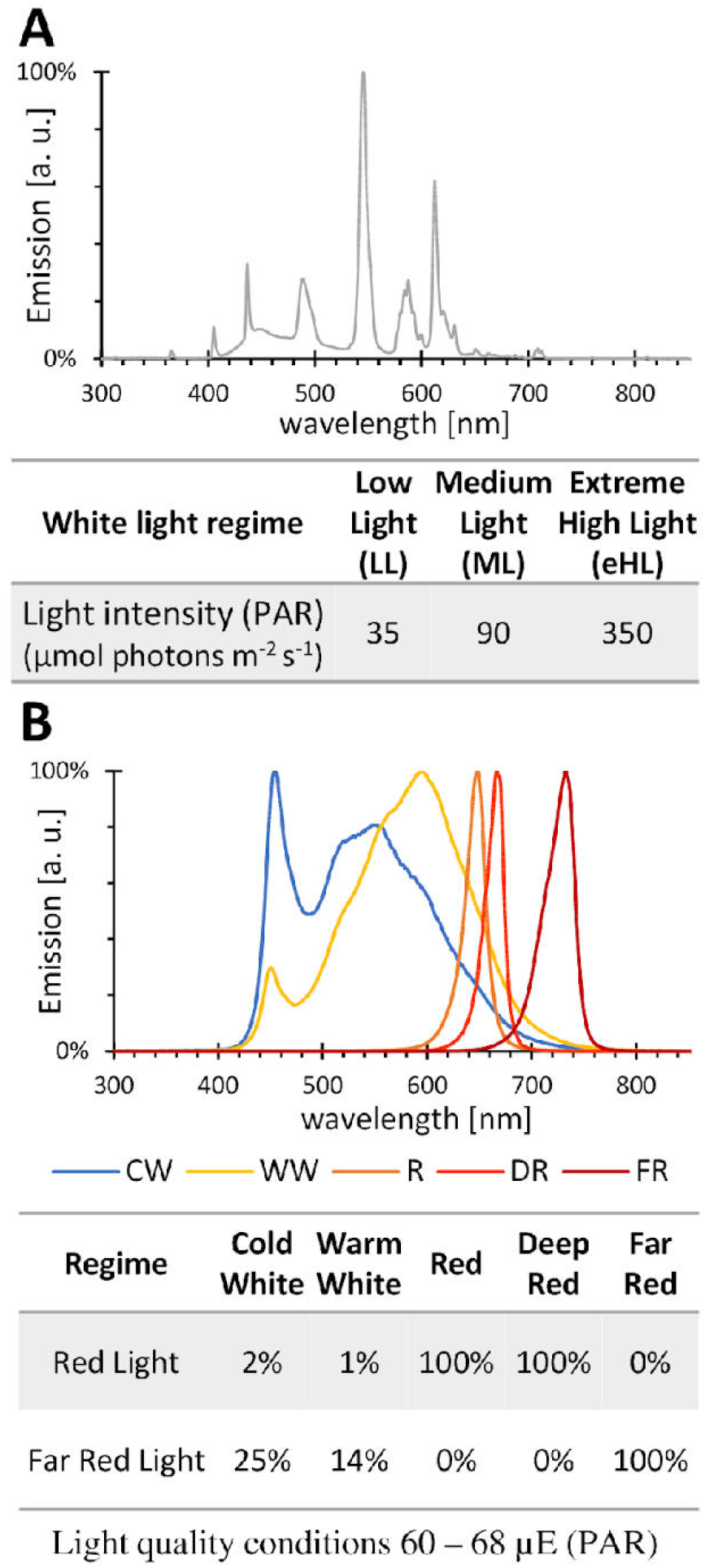
Illumination spectra used in the study. **(A)** Light conditions of the *C. merolae* cell culture grown under various light intensities under white light. The spectra originate from the Panasonic FL40SS/40s ENW37 fluorescent lamps used in the study. Below the light intensity of each culture. **(B)** Spectra of the light sources in the Biogenet™ chamber with a LED array and the table describing the spectral composition. Spectral ranges: cold white light (CWL), 6500K; warm white (WW), 2700K; red light (RL), 630-650 nm; deep red light (DR), 650-670 nm; far red light (FRL), 710-740 nm.

**Supplementary Figure 2.**
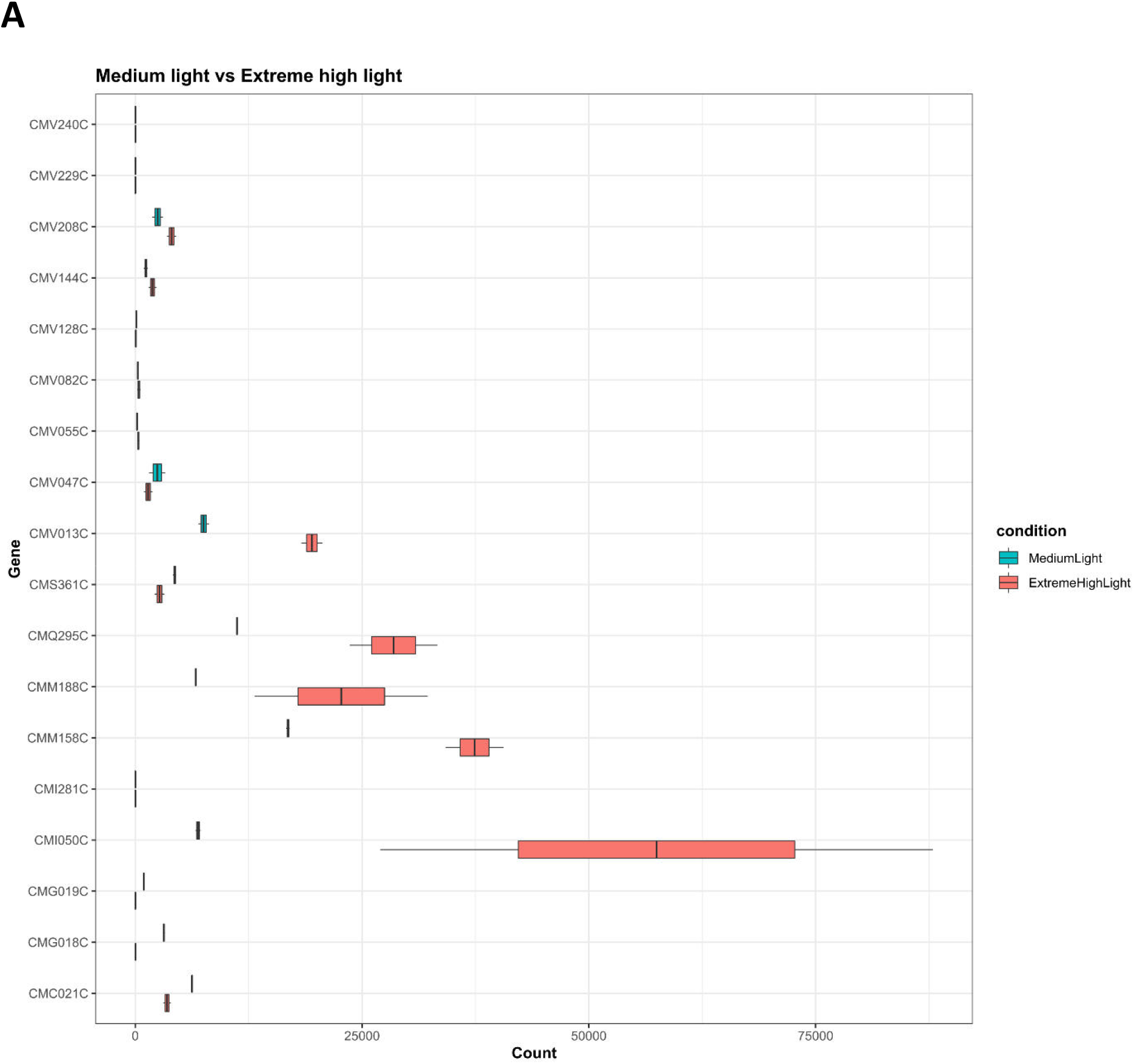
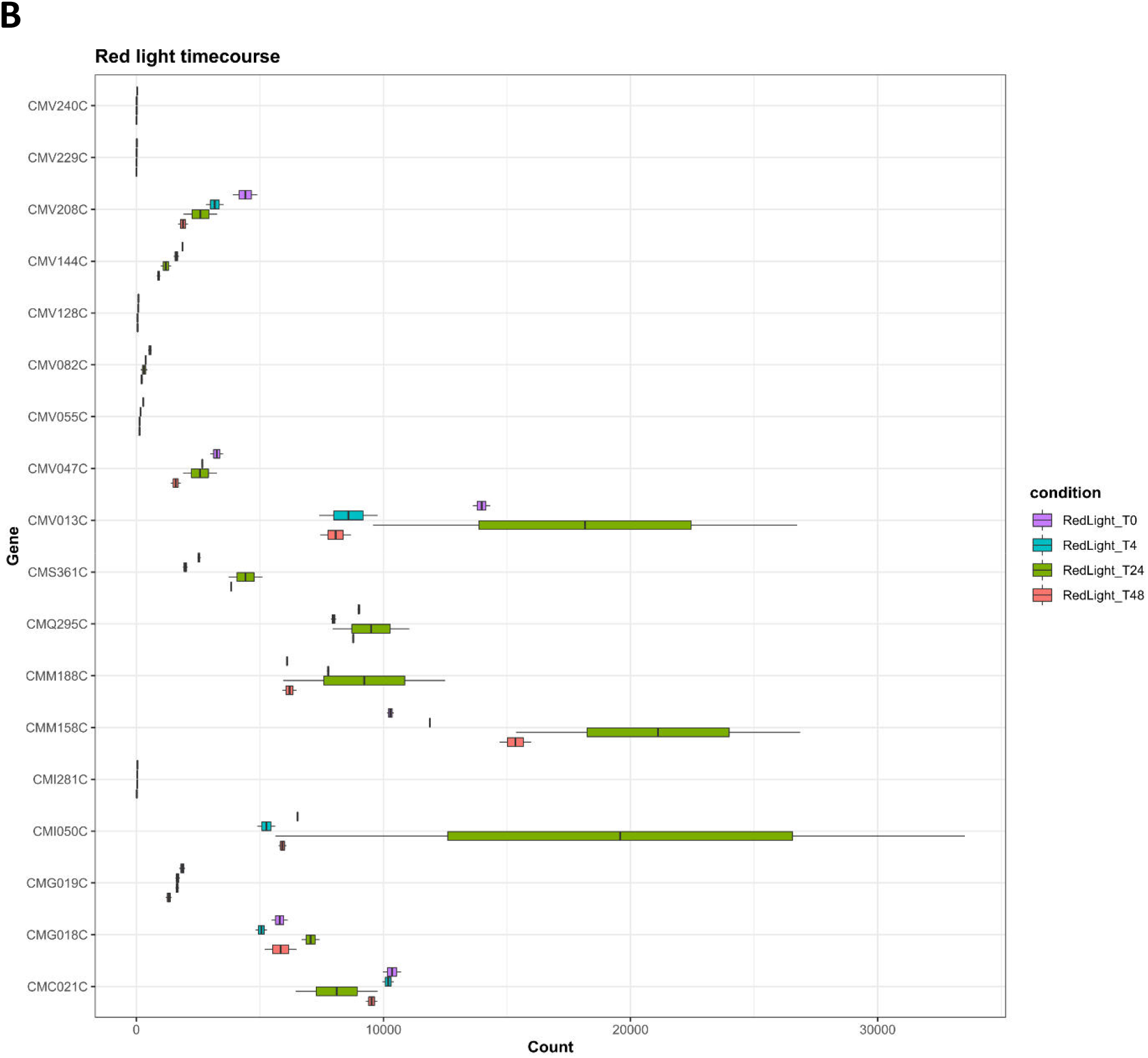
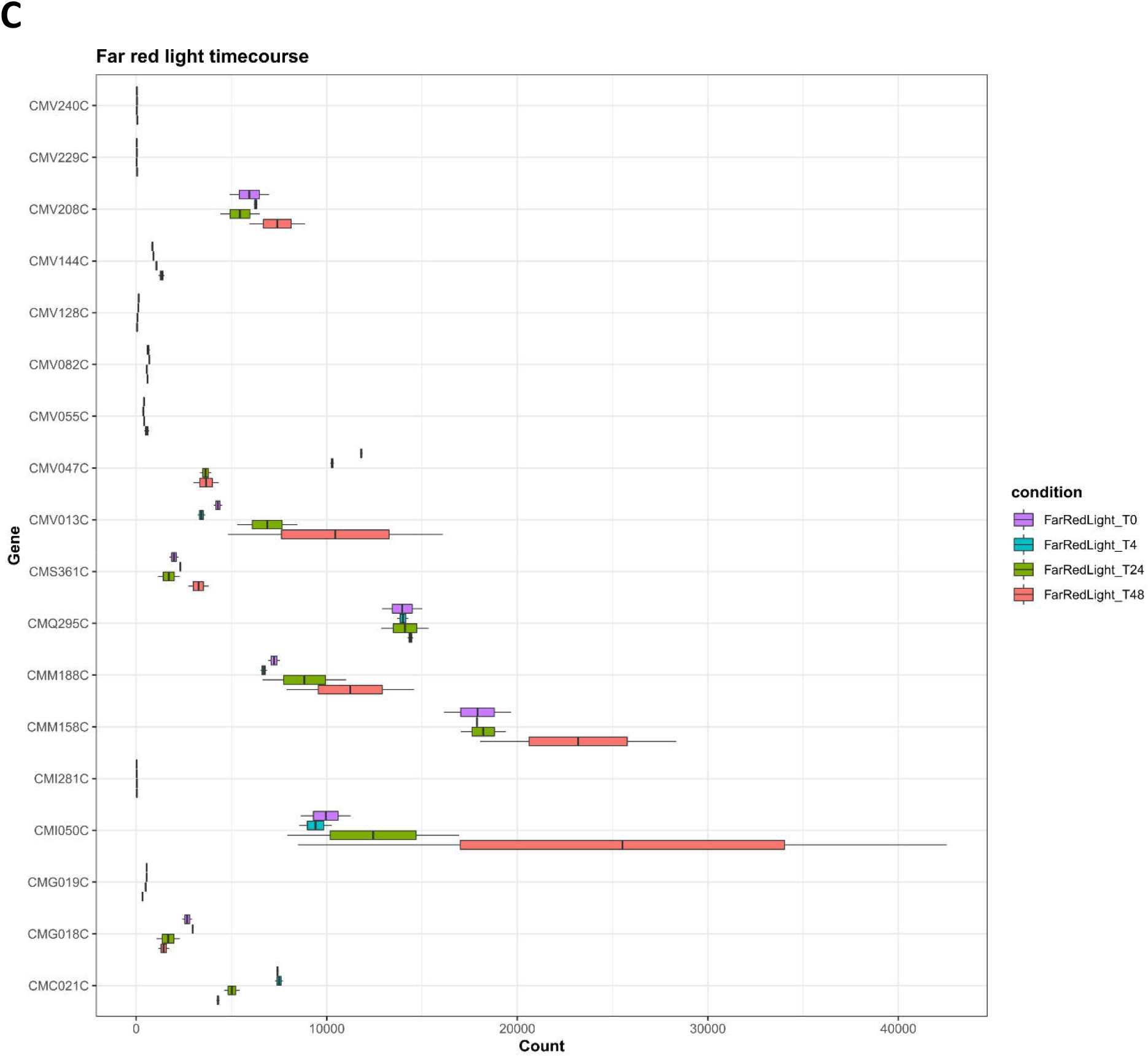
Relative expression of genes of interest. Gene expression counts for all discussed genes as normalized by DESeq2, plotted for comparisons. **(A)** ML versus eHL treatment, **(B)** RL time course and **(C)** FRL time course.

**Supplementary Table 1.** List of genes selected for the transcriptomic analysis.

| **Photosynthetic complex and metabolic pathways** | **Gene accession no.** | **Protein identity and function** |
| --- | --- | --- |
| PSI complex | CMV135C | PsaA (reaction centre) |
|  | CMV136C | PsaB (reaction centre) |
|  | CMV059C | PsaC (stromal subunit, ferredoxin binding pocket) |
|  | CMV144C | PsaD (stromal subunit, ferredoxin binding pocket) |
|  | CMV128C | PsaE (stromal subunit, ferredoxin binding pocket) |
|  | CMV201C | PsaF |
|  | CMV202C | PsaJ |
|  | CMV055C | PsaK (association of LHCI) |
|  | CMV236C | PsaL |
|  | CMP086C | PsaO |
| LHCI antenna | CMN234C | Lhcr |
|  | CMN235C | Lhcr |
|  | CMQ142C | Lhcr |
| PSII complex | CMV047C | PsbA (D1 reaction centre) |
|  | CMV081C | PsbD (D2 reaction centre) |
|  | CMV082C | PsbC (CP43 inner antenna) |
|  | CMV124C | PsbB (CP47 inner antenna) |
|  | CMI290C | PsbO (OEC component) |
|  | CMC133C | PsbQ' (OEC component) |
|  | CMI248C | PsbU (OEC component) |
|  | CMV208C | PsbV (cyt *c*<sub>550</sub>, OEC component) |
| | CMV231C | PsbE (α-cyt *b*<sub>559</sub>) |
| | CMV230C | PsbF (β-cyt *b*<sub>559</sub>) |
|  | CMV127C | PsbH |
|  | CMV240C | PsbI |
|  | CMV140C | PsbK |
|  | CMV229C | PsbL |
|  | CMT182C | PsbM |
|  | CMV089C | PsbW |
|  | CMK176C | Psb27 |
| PBS antenna | CMV063C | PC alpha chain |
|  | CMV064C | PC beta chain |
|  | CMV158C | APC alpha chain |
|  | CMV159C | APC beta chain |
|  | CMV051C | PBS rod-core linker polypeptide |
|  | CMP166C | PC-associated rod linker protein |
| other photosynthetic components and related components | CMV014C | ribulose biphosphate carboxylase small chain (RuBisCO) |
|  | CMV013C | ribulose biphosphate carboxylase large chain (RuBisCO) |
|  | CMV024C | Mg-protoporphyrin IX chelatase (Chl and haem biosynthesis) |
|  | CMJ212C | similar to cytochrome *c* |
|  | CMV096C | PetB cytochrome *b*<sub>6</sub> |
|  | CMV129C | FtsH homolog (thylakoid biogenesis and PSII repair cycle) |
|  | CMQ295C | FtsH (thylakoid biogenesis and PSII repair cycle) |
|  | CMV225C | thylakoid ATP synthase CF1 alpha chain |
|  | CMV196C | thylakoid ATP synthase CF1 beta chain |
|  | CMV222C | thylakoid ATP synthase CF0 B chain (subunit II) |
|  | CMV209C | PetJ cytochrome *c*<sub>6</sub> |
|  | CMG162C | similar to ferredoxin component (biphenyl dioxygenase, Rieske iron-sulfur component related protein) |
|  | CMI281C | cytochrome *b<sub>6</sub>f* complex iron-sulfur subunit precursor |
|  | CMJ096C | thioredoxin (multiple metabolic pathways) |
|  | CMO314C | Photosystem II stability/assembly factor HCF136 |
|  | CMF012C | PPDK pyruvate phosphate dikinase (CO<sub>2</sub> fixation) |
|  | CMN328C | phosphate/phosphoenolpyruvate translocator precursor |
|  | CMS231C | TSPO (ABA-dependent haem binding protein involved in retrograde chloroplast/nucleus transport) |
|  | CMM072C | CmSIG4 nuclear encoded sigma factor 4 (activation of transcription of psbA and ycf17 transcripts) |
|  | CMK044C | CmSIG1 nuclear encoded sigma factor 1 (transcriptional regulation of chloroplast genes in HL) |
|  | CMQ213C | CmSIG2 nuclear encoded sigma factor 2 (RL and FRL-induced signalling between nucleus and chloroplast) |
|  | CMR165C | CmSIG3 nuclear encoded sigma factor 3 (plastid genes transcription) |
| ROS detoxification,<br>blue light sensing,<br>circadian rhythms | CMI050C | catalase |
|  | CMM158C | CmSTAPX stromal ascorbate peroxidase |
|  | CML130C | CmPHR2 cryptochrome DASH (ssDNA damage) |
|  | CMM076C | CmPHR3 cryptochrome DASH (ssDNA damage) |
|  | CMQ453 | CmPHR7 cryptochrome |
|  | CMV057C | ORF193 (putative blue light receptor) |
| respiration | CMT434C | mitochondrial F-type ATPase F1 subunit alpha, precursor |
|  | CMH197C | mitochondrial F-type ATPase F1 subunit beta, precursor |
|  | CMS272C | isocitrate dehydrogenase subunit 2, mitochondrial (Krebs cycle) |
|  | CMP193C | malate dehydrogenase, mitochondrial (Krebs cycle) |
|  | CMN084C | ADP/ATP translocase |
| heat shock proteins | CMJ101C | CmsHSP2 Small Heat Shock Protein 20 |
|  | CMJ100C | CmsHSP1 Small Heat Shock Protein 20 |
|  | CMP145C | heat shock protein Hsp70 |
|  | CMQ224C | heat shock protein of Hsp90 family |
|  | CMS343C | molecular chaperone of Hsp110 family |
|  | CMI251C | heat shock protein ClpB |
|  | CML205C | heat shock protein DnaK |
| cell cycle and<br>organelle division | CMS361C | FtsZ1 (mitochondrial division) |
|  | CMS004C | FtsZ2 (chloroplast division) |
| lipid biosynthesis and<br>metabolism | CMA017C | glycerol-3-phosphate acyltransferase (GPAT) |
|  | CMK217C | acyl-CoA:diacylglycerol acyltransferase (DGAT) |
|  | CMM188C | acetyl-CoA carboxylase (ACC) |
|  | CME186C | long chain acyl-CoA synthetase |
|  | CMK139C | enoyl-CoA hydratase, mitochondrial precursor |
|  | CMR012C | chloroplast UDP-sulfoquinovose synthase |
| nitrogen metabolism | CMJ282C | CmMYB1 activator of the nitrogen assimilation pathway |
|  | CMG018C | nitrate transporter (NRT) |
|  | CMG019C | nitrate reductase (NR) |
|  | CMG021C | nitrite reductase (NIR) |
| glycolysis | CMC021C | pyruvate kinase |
|  | CMK188C | phosphoglycerate mutase |
| other metabolic pathways<br>and proteins of unknown<br>function | CMC045C | 60S ribosomal protein L40, ubiquitin fusion protein |
|  | CMC148C | aspartate aminotransferase (amino acid metabolism) |
|  | CMC180C | putative protein disulfide-isomerase |
|  | CME050C | similar to nuclear LIM interactor-interacting factor |
|  | CMH166C | DNA gyrase subunit B |
|  | CMH226C | eukaryotic translation elongation factor 1 alpha (eEF-1a) |
|  | CMI213C | calnexin |
|  | CMJ260C | similar to prenylated Rab acceptor PRA1 |
|  | CML276C | chaperonin containing TCP1, subunit 5 |
|  | CMM237C | actin (cytoskeleton) |
|  | CMN169C | histone H4 |
|  | CMO083C | similar to processing protease (protein quality control) |
|  | CMS362C | CmNCED epoxycarotenoid dioxygenase (stress tolerance) |
|  | CMP198C | similar to bestrophin (anion channel) |
|  | CMP239C | 14-3-3 protein |
|  | CMQ074C | small GTP-binding protein Arf1 |
|  | CMQ122C | small GTP-binding protein of Rab family, Rab6 |
|  | CMQ225C | 20S core proteasome subunit alpha 5 |
|  | CMQ424C | nuclear poly(A) polymerase |
|  | CMR299C | stress-induced phosphoprotein STI1 |
|  | CMR384C | ubiquitin carboxyl-terminal hydrolase |
|  | CMR457C | histone H2A.Z variant |
|  | CMS243C | DNA gyrase subunit A |
|  | CMT098C | ubiquitinyl hydrolase 1 |
|  | CMT113C | putative phosphoribosyl diphosphate synthase; ribose-phosphate pyrophosphokinase |
|  | CMT173C | ATP-dependent RNA helicase |
|  | CMT197C | 40S ribosomal protein S27 |
|  | CMT305C | argininosuccinate synthase |
|  | CMM134C | uncharacterized protein, similar to histone deacetylase HDT1 |

*Gene accession number according to Matsuzaki et al. (2004) with later updates. Proteins are grouped according to their specific function in the metabolic pathways shown on the left.*

## Literature cited

Abram M, Białek R, Szewczyk S, Karolczak J, Gibasiewicz K, Kargul J (2020) Remodeling of excitation energy transfer in extremophilic red algal PSI-LHCI complex during light adaptation. Biochim Biophys Acta BBA - Bioenerg 1861: 148093

Ago H, Adachi H, Umena Y, Tashiro T, Kawakami K, Kamiya N, Tian L, Han G, Kuang T, Liu Z, et al (2016) Novel Features of Eukaryotic Photosystem II Revealed by Its Crystal Structure Analysis from a Red Alga. J Biol Chem 291: 5676–5687

Allen JF, de Paula WBM, Puthiyaveetil S, Nield J (2011) A structural phylogenetic map for chloroplast photosynthesis. Trends Plant Sci 16: 645–655

Anderson JM, Chow WS, Goodchild DJ (1988) Thylakoid Membrane Organisation in Sun/Shade Acclimation. Funct Plant Biol 15: 11–26

Antoshvili M, Caspy I, Hippler M, Nelson N (2019) Structure and function of photosystem I in *Cyanidioschyzon merolae*. Photosynth Res 139: 499–508

Archibald JM, Keeling PJ (2002) Recycled plastids: a ‘green movement’ in eukaryotic evolution. Trends Genet 18: 577–584

Athanasiou K (2007) Kinetics of Acclimation in the Changing Light Environment. The University of Manchester (United Kingdom)

Athanasiou K, Dyson BC, Webster RE, Johnson GN (2010) Dynamic Acclimation of Photosynthesis Increases Plant Fitness in Changing Environments. Plant Physiol 152: 366–373

Bailey S, Walters RG, Jansson S, Horton P (2001) Acclimation of *Arabidopsis thaliana* to the light environment: the existence of separate low light and high light responses. Planta 213: 794–801

Baker NR (2008) Chlorophyll Fluorescence: A Probe of Photosynthesis In Vivo. Annu Rev Plant Biol 59: 89–113

Ballottari M, Dall’Osto L, Morosinotto T, Bassi R (2007) Contrasting Behavior of Higher Plant Photosystem I and II Antenna Systems during Acclimation. J Biol Chem 282: 8947–8958

Bellafiore S, Barneche F, Peltier G, Rochaix J-D (2005) State transitions and light adaptation require chloroplast thylakoid protein kinase STN7. Nature 433: 892–895

Black CS, Garside EL, MacMillan AM, Rader SD (2016) Conserved structure of Snu13 from the highly reduced spliceosome of *Cyanidioschyzon merolae*. Protein Sci 25: 911–916

Boardman NK (1977) Comparative Photosynthesis of Sun and Shade Plants. Annu Rev Plant Physiol 28: 355–377

Bolychevtseva YV, Tropin IV, Stadnichuk IN (2021) State 1 and State 2 in Photosynthetic Apparatus of Red Microalgae and Cyanobacteria. Biochem Mosc 86: 1181–1191

Bos I, Bland KM, Tian L, Croce R, Frankel LK, van A, Bricker TM, Wientjes E (2017) Multiple LHCII antennae can transfer energy efficiently to a single Photosystem I. Biochim Biophys Acta - Bioenerg 1858: 371–378

Burger G, Saint-Louis D, Gray MW, Lang BF (1999) Complete Sequence of the Mitochondrial DNA of the Red Alga *Porphyra purpurea:* Cyanobacterial Introns and Shared Ancestry of Red and Green Algae. Plant Cell 11: 1675–1694

Burki F, Kaplan M, Tikhonenkov DV, Zlatogursky V, Minh BQ, Radaykina LV, Smirnov A, Mylnikov AP, Keeling PJ (2016) Untangling the early diversification of eukaryotes: a phylogenomic study of the evolutionary origins of Centrohelida, Haptophyta and Cryptista. Proc R Soc B Biol Sci 283: 20152802

Busch A, Nield J, Hippler M (2010) The composition and structure of photosystem I-associated antenna from *Cyanidioschyzon merolae*. Plant J 62: 886–897

Calzadilla PI, Kirilovsky D (2020) Revisiting cyanobacterial state transitions. Photochem Photobiol Sci 19: 585–603

Canonico M, Konert G, Crepin A, Šedivá B, Kaňa R (2021) Gradual Response of Cyanobacterial Thylakoids to Acute High-Light Stress—Importance of Carotenoid Accumulation. Cells 10: 1916

Canonico M, Konert G, Kaňa R (2020) Plasticity of Cyanobacterial Thylakoid Microdomains Under Variable Light Conditions. Front Plant Sci. doi: 10.3389/fpls.2020.586543

Campbell D, Hurry V, Clarke AK, Gustafsson P, Öquist G (1998) Chlorophyll Fluorescence Analysis of Cyanobacterial Photosynthesis and Acclimation. Microbiol Mol Biol Rev 62: 667–683

Chang L, Tian L, Ma F, Mao Z, Liu X, Han G, Wang W, Yang Y, Kuang T, Pan J, et al (2020) Regulation of photosystem I-light-harvesting complex I from a red alga *Cyanidioschyzon merolae* in response to light intensities. Photosynth Res 146: 287–297

Chen S, Zhou Y, Chen Y, Gu J (2018) fastp: an ultra-fast all-in-one FASTQ preprocessor. Bioinformatics 34: i884–i890

Ciniglia C, Yoon HS, Pollio A, Pinto G, Bhattacharya D (2004) Hidden biodiversity of the extremophilic Cyanidiales red algae. Mol Ecol 13: 1827–1838

Croce R (2020) Beyond ‘seeing is believing’: the antenna size of the photosystems in vivo. New Phytol. doi: 10.1111/nph.16758

Cunningham FX, Dennenberg RJ, Mustardy L, Jursinic PA, Gantt E (1989) Stoichiometry of Photosystem I, Photosystem II, and Phycobilisomes in the Red Alga *Porphyridium cruentum* as a Function of Growth Irradiance. Plant Physiol 91: 1179–1187

Cunningham FX, Lee H, Gantt E (2007) Carotenoid Biosynthesis in the Primitive Red Alga *Cyanidioschyzon merolae*. Eukaryot Cell 6: 533–545

Delphin E, Duval J-C, Kirilovsky D (1995) Comparison of state 1-state 2 transitions in the green alga *Chlamydomonas reinhardtii* and in the red alga *Rhodella violacea:* effect of kinase and phosphatase inhibitors. Biochim Biophys Acta BBA - Bioenerg 1232: 91–95

Delphin E, Duval J-C, Etienne A-L, Kirilovsky D (1996) State Transitions or ΔpH-Dependent Quenching of Photosystem II Fluorescence in Red Algae. Biochemistry 35: 9435–9445

Delphin E, Duval J-C, Etienne A-L, Kirilovsky D (1998) ΔpH-Dependent Photosystem II Fluorescence Quenching Induced by Saturating, Multiturnover Pulses in Red Algae. Plant Physiol 118: 103–113

Depège N, Bellafiore S, Rochaix J-D (2003) Role of Chloroplast Protein Kinase Stt7 in LHCII Phosphorylation and State Transition in *Chlamydomonas*. Science 299: 1572–1575

Dietzel L, Bräutigam K, Pfannschmidt T (2008) Photosynthetic acclimation: State transitions and adjustment of photosystem stoichiometry – functional relationships between short-term and long-term light quality acclimation in plants. FEBS J 275: 1080–1088

Dobin A, Davis CA, Schlesinger F, Drenkow J, Zaleski C, Jha S, Batut P, Chaisson M, Gingeras TR (2013) STAR: ultrafast universal RNA-seq aligner. Bioinformatics 29: 15–21

Dominguez-Martin MA, Sauer PV, Sutter M, Kirst H, Bina D, Greber BJ, Nogales E, Polívka T, Kerfeld CA (2021) Structure of the Quenched Cyanobacterial OCP-Phycobilisome Complex. 2021.11.15.468719

Duanmu D, Rockwell NC, Lagarias JC (2017) Algal light sensing and photoacclimation in aquatic environments. Plant Cell Environ 40: 2558–2570

Enami I, Kikuchi S, Fukuda T, Ohta H, Shen J-R (1998) Binding and Functional Properties of Four Extrinsic Proteins of Photosystem II from a Red Alga, *Cyanidium caldarium*, As Studied by Release-Reconstitution Experiments. Biochemistry 37: 2787–2793

Enami I, Okumura A, Nagao R, Suzuki T, Iwai M, Shen J-R (2008) Structures and functions of the extrinsic proteins of photosystem II from different species. Photosynth Res 98: 349–363

Engelmann E, Zucchelli G, Casazza AP, Brogioli D, Garlaschi FM, Jennings RC (2006) Influence of the Photosystem I-Light Harvesting Complex I Antenna Domains on Fluorescence Decay. Biochemistry 45: 6947–6955

Flannery SE, Hepworth C, Wood WHJ, Pastorelli F, Hunter CN, Dickman MJ, Jackson PJ, Johnson MP (2021) Developmental acclimation of the thylakoid proteome to light intensity in *Arabidopsis*. Plant J 105: 223–244

Foyer CH, Noctor G (2005) Redox Homeostasis and Antioxidant Signaling: A Metabolic Interface between Stress Perception and Physiological Responses. Plant Cell 17: 1866–1875

Fujita Y, Murakami A, Aizawa K, Ohki K (1994) Short-term and Long-term Adaptation of the Photosynthetic Apparatus: Homeostatic Properties of Thylakoids. In DA Bryant, ed, Mol. Biol. Cyanobacteria. Springer Netherlands, Dordrecht, pp 677–692

Fujiwara T, Kanesaki Y, Hirooka S, Era A, Sumiya N, Yoshikawa H, Tanaka K, Miyagishima S-Y (2015) A nitrogen source-dependent inducible and repressible gene expression system in the red alga *Cyanidioschyzon merolae*. Front Plant Sci. doi: 10.3389/fpls.2015.00657

Garside EL, Whelan TA, Stark MR, Rader SD, Fast NM, MacMillan AM (2019) Prp8 in a Reduced Spliceosome Lacks a Conserved Toggle that Correlates with Splicing Complexity across Diverse Taxa. J Mol Biol 431: 2543–2553

Gibasiewicz K, Szrajner A, Ihalainen JA, Germano M, Dekker JP, van Grondelle R (2005) Characterization of Low-Energy Chlorophylls in the PSI-LHCI Supercomplex from *Chlamydomonas reinhardtii*. A Site-Selective Fluorescence Study. J Phys Chem B 109: 21180–21186

Gobets B, van Grondelle R (2001) Energy transfer and trapping in photosystem I. Biochim Biophys Acta 1507: 80–99

Goss R, Lepetit B (2015) Biodiversity of NPQ. J Plant Physiol 172: 13–32

Gray GR, Savitch LV, Ivanov AG, Huner NPA (1996) Photosystem II Excitation Pressure and Development of Resistance to Photoinhibition (II. Adjustment of Photosynthetic Capacity in Winter Wheat and Winter Rye). Plant Physiol 110: 61–71

Hager A (1967) Studies on the light-induced reversible xanthophyll-conversions in Chlorella and spinach leaves. Planta 74: 148–172

Haniewicz P, Abram M, Nosek L, Kirkpatrick J, El-Mohsnawy E, Olmos JDJ, Kouřil R, Kargul JM (2018) Molecular Mechanisms of Photoadaptation of Photosystem I Supercomplex from an Evolutionary Cyanobacterial/Algal Intermediate. Plant Physiol 176:1433–1451

Horton P, Johnson MP, Perez-Bueno ML, Kiss AZ, Ruban AV (2008) Photosynthetic acclimation: Does the dynamic structure and macro-organisation of photosystem II in higher plant grana membranes regulate light harvesting states? FEBS J 275: 1069–1079

Ifuku K (2015) Localization and functional characterization of the extrinsic subunits of photosystem II: an update. Biosci Biotechnol Biochem 79: 1223–1231

Ihalainen JA, Jensen PE, Haldrup A, van Stokkum IHM, van Grondelle R, Scheller HV, Dekker JP (2002) Pigment Organization and Energy Transfer Dynamics in Isolated Photosystem I (PSI) Complexes from *Arabidopsis thaliana* Depleted of the PSI-G, PSI-K, PSI-L, or PSI-N Subunit. Biophys J 83: 2190–2201

Itoh R, Takano H, Ohta N, Miyagishima S, Kuroiwa H, Kuroiwa T (1999) Two ftsH-family genes encoded in the nuclear and chloroplast genomes of the primitive red alga *Cyanidioschyzon merolae*. Plant Mol Biol 41: 321–337

Jennings RC, Zucchelli G, Croce R, Garlaschi FM (2003) The photochemical trapping rate from red spectral states in PSI–LHCI is determined by thermal activation of energy transfer to bulk chlorophylls. Biochim Biophys Acta 1557: 91–98

Kaňa R, Kotabová E, Komárek O, Šedivá B, Papageorgiou GC, Govindjee, Prášil O (2012) The slow S to M fluorescence rise in cyanobacteria is due to a state 2 to state 1 transition. Biochim Biophys Acta BBA - Bioenerg 1817: 1237–1247

Kaňa R, Kotabová E, Lukeš M, Papáček S, Matonoha C, Liu L-N, Prášil O, Mullineaux CW (2014) Phycobilisome Mobility and Its Role in the Regulation of Light Harvesting in Red Algae. Plant Physiol 165: 1618–1631

Kargul J, Barber J (2008) Photosynthetic acclimation: Structural reorganisation of light harvesting antenna – role of redox-dependent phosphorylation of major and minor chlorophyll a/b binding proteins. FEBS J 275: 1056–1068

Kerfeld CA, Sawaya MR, Brahmandam V, Cascio D, Ho KK, Trevithick-Sutton CC, Krogmann DW, Yeates TO (2003) The Crystal Structure of a Cyanobacterial Water-Soluble Carotenoid Binding Protein. Structure 11: 55–65

Kirilovsky D (2007) Photoprotection in cyanobacteria: the orange carotenoid protein (OCP)-related non-photochemical-quenching mechanism. Photosynth Res 93: 7

Kirilovsky D, Kaňa R, Prášil O (2014) Mechanisms Modulating Energy Arriving at Reaction Centers in Cyanobacteria. In B Demmig-Adams, G Garab, W Adams III, Govindjee, eds, Non-Photochem. Quenching Energy Dissipation Plants Algae Cyanobacteria. Springer Netherlands, Dordrecht, pp 471–501

Kirst H, Formighieri C, Melis A (2014) Maximizing photosynthetic efficiency and culture productivity in cyanobacteria upon minimizing the phycobilisome light-harvesting antenna size. Biochim Biophys Acta BBA - Bioenerg 1837: 1653–1664

Konert G, Steinbach G, Canonico M, Kaňa R (2019) Protein arrangement factor: a new photosynthetic parameter characterizing the organization of thylakoid membrane proteins. Physiol Plant 166: 264–277

Kosuge K, Tokutsu R, Kim E, Akimoto S, Yokono M, Ueno Y, Minagawa J (2018) LHCSR1-dependent fluorescence quenching is mediated by excitation energy transfer from LHCII to photosystem I in *Chlamydomonas reinhardtii*. Proc Natl Acad Sci 115: 3722–3727

Krupnik T, Kotabová E, Bezouwen LS van, Mazur R, Garstka M, Nixon PJ, Barber J, Kaňa R, Boekema EJ, Kargul J (2013) A Reaction Center-dependent Photoprotection Mechanism in a Highly Robust Photosystem II from an Extremophilic Red Alga, *Cyanidioschyzon merolae*. J Biol Chem 288: 23529–23542

Kuroiwa T (1998) The primitive red algae *Cyanidium caldarium* and *Cyanidioschyzon merolae* as model system for investigating the dividing apparatus of mitochondria and plastids. BioEssays 20: 344–354

Li H, Handsaker B, Wysoker A, Fennell T, Ruan J, Homer N, Marth G, Abecasis G, Durbin R, 1000 Genome Project Data Processing Subgroup (2009a) The Sequence Alignment/Map format and SAMtools. Bioinformatics 25: 2078–2079

Li X-P, Müller-Moulé P, Gilmore AM, Niyogi KK (2002) PsbS-dependent enhancement of feedback de-excitation protects photosystem II from photoinhibition. Proc Natl Acad Sci 99:15222–15227

Li Z, Wakao S, Fischer BB, Niyogi KK (2009b) Sensing and responding to excess light. Annu Rev Plant Biol 60: 239–260

Liu L-N (2016) Distribution and dynamics of electron transport complexes in cyanobacterial thylakoid membranes. Biochim Biophys Acta BBA - Bioenerg 1857: 256–265

Liu L-N, Bryan SJ, Huang F, Yu J, Nixon PJ, Rich PR, Mullineaux CW (2012) Control of electron transport routes through redox-regulated redistribution of respiratory complexes. Proc Natl Acad Sci 109: 11431–11436

Love MI, Huber W, Anders S (2014) Moderated estimation of fold change and dispersion for RNA-seq data with DESeq2. Genome Biol 15: 550

Luca PD, Taddei R, Varano L (1978) *Cyanidioschyzon merolae:* a new alga of thermal acidic environments. Webbia 33: 37–44

Ma M, Liu Y, Bai C, Yong JWH (2021) The Significance of Chloroplast NAD(P)H Dehydrogenase Complex and Its Dependent Cyclic Electron Transport in Photosynthesis. Front Plant Sci 12: 661863

MacColl R (1998) Cyanobacterial Phycobilisomes. J Struct Biol 124: 311–334

Matsuzaki M, Misumi O, Shin-i T, Maruyama S, Takahara M, Miyagishima S, Mori T, Nishida K, Yagisawa F, Nishida K, et al (2004) Genome sequence of the ultrasmall unicellular red alga *Cyanidioschyzon merolae* 10D. Nature 428: 653–657

Maxwell K, Johnson GN (2000) Chlorophyll fluorescence—a practical guide. J Exp Bot 51: 659–668

McConnell MD, Koop R, Vasil’ev S, Bruce D (2002) Regulation of the Distribution of Chlorophyll and Phycobilin-Absorbed Excitation Energy in Cyanobacteria. A Structure-Based Model for the Light State Transition. Plant Physiol 130: 1201–1212

Melkozernov AN, Kargul J, Lin S, Barber J, Blankenship RE (2005) Spectral and kinetic analysis of the energy coupling in the PS I-LHC I supercomplex from the green alga *Chlamydomonas reinhardtii* at 77 K. Photosynth Res 86: 203–215

Miller MAE, O’Cualain R, Selley J, Knight D, Karim MF, Hubbard SJ, Johnson GN (2017) Dynamic Acclimation to High Light in *Arabidopsis thaliana* Involves Widespread Reengineering of the Leaf Proteome. Front Plant Sci 8: 1239

Minagawa J (2011) State transitions—The molecular remodeling of photosynthetic supercomplexes that controls energy flow in the chloroplast. Biochim Biophys Acta BBA - Bioenerg 1807: 897–905

Minoda A, Sakagami R, Yagisawa F, Kuroiwa T, Tanaka K (2004) Improvement of culture conditions and evidence for nuclear transformation by homologous recombination in a red alga, *Cyanidioschyzon merolae* 10D. Plant Cell Physiol 45: 667–671

Miyagishima S-Y, Tanaka K (2021) The Unicellular Red Alga *Cyanidioschyzon merolae*— The Simplest Model of a Photosynthetic Eukaryote. Plant Cell Physiol 62: 926–941

Moreira D, Le Guyader H, Philippe H (2000) The origin of red algae and the evolution of chloroplasts. Nature 405: 69–72

Mouget J-L, Tremblin G (2002) Suitability of the Fluorescence Monitoring System (FMS, Hansatech) for measurement of photosynthetic characteristics in algae. Aquat Bot 74: 219–231

Mullineaux CW, Emlyn-Jones D (2005) State transitions: an example of acclimation to low-light stress. J Exp Bot 56: 389–393

Murchie EH, Ruban AV (2020) Dynamic non-photochemical quenching in plants: from molecular mechanism to productivity. Plant J 101: 885–896

Nilsson H, Krupnik T, Kargul J, Messinger J (2014) Substrate water exchange in photosystem II core complexes of the extremophilic red alga *Cyanidioschyzon merolae*. Biochim Biophys Acta 1837: 1257–1262

Nowack ECM, Melkonian M, Glöckner G (2008) Chromatophore Genome Sequence of *Paulinella* Sheds Light on Acquisition of Photosynthesis by Eukaryotes. Curr Biol 18: 410–418

Nozaki H, Takano H, Misumi O, Terasawa K, Matsuzaki M, Maruyama S, Nishida K, Yagisawa F, Yoshida Y, Fujiwara T, et al (2007) A 100%-complete sequence reveals unusually simple genomic features in the hot-spring red alga *Cyanidioschyzon merolae*. BMC Biol 5: 1–8

Ohashi S, Iemura T, Okada N, Itoh S, Furukawa H, Okuda M, Ohnishi-Kameyama M, Ogawa T, Miyashita H, Watanabe T, et al (2010) An overview on chlorophylls and quinones in the photosystem I-type reaction centers. Photosynth Res 104: 305–319

Papageorgiou GC, Tsimilli-Michael M, Stamatakis K (2007) The fast and slow kinetics of chlorophyll a fluorescence induction in plants, algae and cyanobacteria: a viewpoint. Photosynth Res 94: 275–290

Pesaresi P, Scharfenberg M, Weigel M, Granlund I, Schröder WP, Finazzi G, Rappaport F, Masiero S, Furini A, Jahns P, et al (2009) Mutants, Overexpressors, and Interactors of *Arabidopsis* Plastocyanin Isoforms: Revised Roles of Plastocyanin in Photosynthetic Electron Flow and Thylakoid Redox State. Mol Plant 2: 236–248

Pham LV, Janna Olmos JD, Chernev P, Kargul J, Messinger J (2019) Unequal misses during the flash-induced advancement of photosystem II: effects of the S state and acceptor side cycles. Photosynth Res 139: 93–106

Pi X, Tian L, Dai H-E, Qin X, Cheng L, Kuang T, Sui S-F, Shen J-R (2018) Unique organization of photosystem I-light-harvesting supercomplex revealed by cryo-EM from a red alga. Proc Natl Acad Sci U S A 115: 4423–4428

Pribil M, Pesaresi P, Hertle A, Barbato R, Leister D (2010) Role of Plastid Protein Phosphatase TAP38 in LHCII Dephosphorylation and Thylakoid Electron Flow. PLOS Biology. doi: 10.1371/journal.pbio.1000288

Porra RJ, Thompson WA, Kriedemann PE (1989) Determination of accurate extinction coefficients and simultaneous equations for assaying chlorophylls a and b extracted with four different solvents: verification of the concentration of chlorophyll standards by atomic absorption spectroscopy. Biochimica et Biophysica Acta 975: 384–394

Ragan MA, Gutell RR (1995) Are red algae plants? Bot J Linn Soc 118: 81–105

Rakhimberdieva MG, Boichenko VA, Karapetyan NV, Stadnichuk IN (2001) Interaction of phycobilisomes with photosystem II dimers and photosystem I monomers and trimers in the cyanobacterium *Spirulina platensis*. Biochemistry 40: 15780–15788

Reimer KA, Stark MR, Aguilar L-C, Stark SR, Burke RD, Moore J, Fahlman RP, Yip CK, Kuroiwa H, Oeffînger M, et al (2017) The sole LSm complex in *Cyanidioschyzon merolae* associates with pre-mRNA splicing and mRNA degradation factors. RNA 23: 952–967

Sanchez-Tarre V, Kiparissides A (2021) The effects of illumination and trophic strategy on gene expression in *Chlamydomonas reinhardtii*. Algal Res 54: 102186

Sauer PV, Dominguez-Martin MA, Kirst H, Sutter M, Bina D, Greber BJ, Nogales E, Polívka T, Kerfeld CA (2021) Structures of the Cyanobacterial Phycobilisome. 2021.11.15.468712

Schneider CA, Rasband WS, Eliceiri KW (2012) NIH Image to ImageJ: 25 years of image analysis. Nat Methods 9: 671–675

Schumann T, Paul S, Melzer M, Dörmann P, Jahns P (2017) Plant Growth under Natural Light Conditions Provides Highly Flexible Short-Term Acclimation Properties toward High Light Stress. Front Plant Sci 8: 681

Seckbach J (1994) The natural history of Cyanidium (Geitler 1933): past and present perspectives. In J Seckbach, ed, Evolutionary Pathways and Enigmatic Algae: Cyanidium caldarium (Rhodophyta) and Related Cells. Springer Netherlands, Dordrecht, pp 99–112

Shapiguzov A, Ingelsson B, Samol I, Andres C, Kessler F, Rochaix J-D, Vener AV, Goldschmidt-Clermont M (2010) The PPH1 phosphatase is specifically involved in LHCII dephosphorylation and state transitions in *Arabidopsis*. Proc Natl Acad Sci 107: 4782–4787

Snellenburg JJ, Dekker JP, van Grondelle R, van Stokkum IHM (2013) Functional Compartmental Modeling of the Photosystems in the Thylakoid Membrane at 77 K. J Phys Chem B 117: 11363–11371

Sonani RR, Gardiner A, Rastogi RP, Cogdell R, Robert B, Madamwar D (2018) Site, trigger, quenching mechanism and recovery of non-photochemical quenching in cyanobacteria: recent updates. Photosynth Res 137: 171–180

Stark MR, Dunn EA, Dunn WSC, Grisdale CJ, Daniele AR, Halstead MRG, Fast NM, Rader SD (2015) Dramatically reduced spliceosome in *Cyanidioschyzon merolae*. Proc Natl Acad Sci 112: E1191–E1200

Steinbach G, Schubert F, Kaňa R (2015) Cryo-imaging of photosystems and phycobilisomes in *Anabaena* sp. PCC 7120 cells. J Photochem Photobiol B 152: 395–399

Stephens M (2017) False discovery rates: a new deal. Biostatistics 18: 275–294

Strašková A, Steinbach G, Konert G, Kotabová E, Komenda J, Tichý M, Kaňa R (2019) Pigment-protein complexes are organized into stable microdomains in cyanobacterial thylakoids. Biochim Biophys Acta BBA - Bioenerg 1860: 148053

Strassert JFH, Irisarri I, Williams TA, Burki F (2021) A molecular timescale for eukaryote evolution with implications for the origin of red algal-derived plastids. Nat Commun 12: 1879

Tian L, Liu Z, Wang F, Shen L, Chen J, Chang L, Zhao S, Han G, Wang W, Kuang T, et al (2017) Isolation and characterization of PSI–LHCI super-complex and their sub-complexes from a red alga *Cyanidioschyzon merolae*. Photosynth Res 133: 201–214

Tikkanen M, Piippo M, Suorsa M, Sirpiö S, Mulo P, Vainonen J, Vener AV, Allahverdiyeva Y, Aro E-M (2006) State transitions revisited—a buffering system for dynamic low light acclimation of *Arabidopsis*. Plant Mol Biol 62: 795–795

Ueno Y, Aikawa S, Kondo A, Akimoto S (2015) Light adaptation of the unicellular red alga, *Cyanidioschyzon merolae*, probed by time-resolved fluorescence spectroscopy. Photosynth Res 125: 211–218

Ueno Y, Aikawa S, Niwa K, Abe T, Murakami A, Kondo A, Akimoto S (2017) Variety in excitation energy transfer processes from phycobilisomes to photosystems I and II. Photosynth Res 133: 235–243

Vialet-Chabrand S, Matthews JSA, Simkin AJ, Raines CA, Lawson T (2017) Importance of Fluctuations in Light on Plant Photosynthetic Acclimation. Plant Physiol 173: 2163–2179

Walters RG, Horton P (1995) Acclimation of *Arabidopsis thaliana* to the light environment: regulation of chloroplast composition. Planta 197: 475–481

Ware MA, Belgio E, Ruban AV (2015) Photoprotective capacity of non-photochemical quenching in plants acclimated to different light intensities. Photosynth Res 126: 261–274

Wei X, Guo J, Li M, Liu Z (2015) Structural Mechanism Underlying the Specific Recognition between the *Arabidopsis* State-Transition Phosphatase TAP38/PPH1 and Phosphorylated Light-Harvesting Complex Protein Lhcb1. Plant Cell 27: 1113–1127

Wientjes E, van Amerongen H, Croce R (2013a) LHCII is an antenna of both photosystems after long-term acclimation. Biochim Biophys Acta BBA - Bioenerg 1827: 420–426

Wientjes E, van Amerongen H, Croce R (2013b) Quantum Yield of Charge Separation in Photosystem II: Functional Effect of Changes in the Antenna Size upon Light Acclimation. J Phys Chem B 117: 11200–11208

Yagisawa F, Nishida K, Kuroiwa H, Nagata T, Kuroiwa T (2007) Identification and mitotic partitioning strategies of vacuoles in the unicellular red alga *Cyanidioschyzon merolae*. Planta 226: 1017–1029

Yagisawa F, Nishida K, Yoshida M, Ohnuma M, Shimada T, Fujiwara T, Yoshida Y, Misumi O, Kuroiwa H, Kuroiwa T (2009) Identification of novel proteins in isolated polyphosphate vacuoles in the primitive red alga *Cyanidioschyzon merolae*. Plant J 60: 882–893

Yamamoto HY, Nakayama TOM, Chichester CO (1962) Studies on the light and dark interconversions of leaf xanthophylls. Arch Biochem Biophys 97: 168–173

Yamanaka G, Glazer AN, Williams RC (1980) Molecular architecture of a light-harvesting antenna. Comparison of wild type and mutant *Synechococcus* 6301 phycobilisomes. J Biol Chem 255: 11004–11010

Yin Z-H, Johnson GN (2000) Photosynthetic acclimation of higher plants to growth in fluctuating light environments. Photosynth Res 63: 97–107

Yokono M, Murakami A, Akimoto S (2011) Excitation energy transfer between photosystem II and photosystem I in red algae: Larger amounts of phycobilisome enhance spillover. Biochim Biophys Acta 1807: 847–853

Zhang J, Ma J, Liu D, Qin S, Sun S, Zhao J, Sui S-F (2017) Structure of phycobilisome from the red alga *Griffithsia pacifica*. Nature 551: 57–63

Zlenko DV, Galochkina TV, Krasilnikov PM, Stadnichuk IN (2017) Coupled rows of PBS cores and PSII dimers in cyanobacteria: symmetry and structure. Photosynth Res 1–16

